# Goal-directed visual information processing with GABAergic inhibition in parietal cortex

**DOI:** 10.1101/2025.03.26.645504

**Authors:** Zhiyan Wang, Sinah Wiborg, Antonia Wittmann, Nina Beck, Susanna Hirschle, Dominik Aschenbrenner, Markus Becker, Sebastian M. Frank

## Abstract

Goal-directed visual information processing involves tracking relevant visual signals (targets) over space and time. However, goal-irrelevant visual signals (distractors) can interfere with the tracking of targets. The neural mechanisms, which promote the tracking of targets in the challenge of interference from distractors remain elusive. Here, we used time-resolved functional magnetic resonance spectroscopy (fMRS) to measure concentrations of γ-aminobutyric acid (GABA), a chief inhibitory neurotransmitter, and glutamate, a chief excitatory neurotransmitter, in parietal and visual cortex while participants performed a visual tracking task for targets among distractors. We found that the more targets had to be tracked, the greater the concentrations of GABA and glutamate in parietal cortex. In visual cortex only the concentration of glutamate increased with the number of targets. Concentration changes of GABA and glutamate only in parietal cortex were differentially associated with tracking performance: Better target tracking was associated with greater increase in GABA concentration and smaller increase in glutamate concentration. The results of control experiments showed that this differential association between tracking performance and metabolite changes in parietal cortex reflected individual differences in distractor suppression: participants with better distractor suppression during tracking tended to have greater increase in GABA concentration and smaller increase in glutamate concentration. This suggests that target-distractor interference during tracking is minimized by GABAergic suppression of goal-irrelevant distractors in parietal cortex, thereby promoting goal-directed visual information processing.

## Introduction

Tracking visual signals relevant to current goals across space and time is critical for adaptive behavior (Carrasco, 2011; Cavanagh & Alvarez, 2005; Chun et al., 2011; Egeth & Yantis, 1997; Gazzaley & Nobre, 2012; Scholl, 2001; Van Ede & Nobre, 2023; Xu & Chun, 2009). However, in a dynamically changing environment goal-irrelevant distractor signals compete for processing and can interfere with tracking of goal-relevant target signals. How is the interference between targets and distractors resolved? Previous results showed that there are distinct neural representations of targets and distractors in visual cortex (Christophel et al., 2012; Ester et al., 2013; Harrison & Tong, 2009; Rademaker et al., 2019; Serences, 2016; Sprague et al., 2014, 2015) and parietal cortex (Bracci et al., 2017; Buschman & Miller, 2007; Vaziri-Pashkam & Xu, 2017). Especially the representation of targets in parietal cortex was found to be robust against interference from distractors (Bettencourt & Xu, 2016; Lorenc et al., 2018; Xu, 2024). However, it is unclear which neural mechanisms promote the emergence of target representations in parietal cortex that are robust to distractor interference.

A crucial neural mechanism for minimizing target-distractor interference could be inhibitory processing with γ-aminobutyric acid (GABA). GABA is a chief inhibitory neurotransmitter (Isaacson & Scanziani, 2011; Petroff, 2002; Schmidt-Wilcke et al., 2018; Tremblay et al., 2016; Tritsch et al., 2016) and critical for suppressing spontaneous activity related to distraction (Bang et al., 2018; De La Vega et al., 2014; Frangou et al., 2019; Frank et al., 2022; Shibata et al., 2017; H. R. Snyder et al., 2010; Tamaki et al., 2020; Yamada et al., 2024). While reduced GABAergic function can impair the selection of goal-relevant signals (H. R. Snyder et al., 2010), increased GABAergic activity is associated with enhanced efficiency in resolving competitions between task-relevant and task-irrelevant signals (De La Vega et al., 2014). On a cellular level, GABAergic inhibition minimizes interneuron correlations and promotes goal-relevant sensory signal processing (Cohen & Maunsell, 2009; Harris & Thiele, 2011; A. C. Snyder et al., 2016; Srivastava et al., 2023).

Here we investigated whether and, if so, how GABAergic inhibition in parietal cortex minimizes target-distractor interference. We used a recently advanced imaging method referred to as time-resolved functional magnetic resonance spectroscopy (fMRS) (Apšvalka et al., 2015; Craven et al., 2024; Ip & Bridge, 2022; Koolschijn et al., 2023; Lally et al., 2014; Mullins, 2018, 2024; Stanley & Raz, 2018) to measure concentrations of GABA and Glx (corresponding to the composite measure of glutamate and its precursor glutamine) in parietal and visual cortex while participants performed a multiple object tracking (MOT) task (Cavanagh & Alvarez, 2005; Meyerhoff et al., 2017; Pylyshyn & Storm, 1988). In different conditions of MOT participants tracked two or four targets among multiple distractors (henceforth referred to as low and high load conditions), to increase target-distractor interference with increasing load. Importantly, the total number of stimuli (sum of targets and distractors) remained identical across conditions to maintain constant visual input (Alvarez & Franconeri, 2007; Cavanagh & Alvarez, 2005).

We found that GABA and glutamate concentrations in parietal cortex increased in MOT with high compared to low load. Critically, a greater increase in GABA concentration and lower increase in glutamate concentration in parietal cortex predicted better tracking performance in MOT with high load. In visual cortex only the concentration of glutamate increased with load and no association with tracking performance was found.

This suggests that GABAergic inhibition in parietal cortex is crucial for tracking performance under conditions of high target-distractor interference. However, minimizing target-distractor interference in MOT with high load could occur through enhancing targets (Brefczynski & DeYoe, 1999; Rorden, 2008), or suppressing distractors (Serences et al., 2004; Wöstmann et al., 2022), or a combination of both (Föcker et al., 2023). To further elucidate the mechanisms underlying GABAergic inhibition in parietal cortex in MOT with high load, we conducted control experiments in which we separately varied the need to enhance targets and suppress distractors during tracking (Bettencourt & Somers, 2009). The results of the control experiments showed a significant association between parietal GABA and distractor suppression, such that a greater increase in GABA concentration predicted better tracking performance with increasing need to suppress distractors during MOT. No significant association between parietal GABA and target enhancement was found.

Together, the results of this study suggest that GABAergic inhibition in parietal cortex minimizes target-distractor interference through distractor suppression, which may promote the emergence of robust target representations. This effect of GABA was found in parietal but not in visual cortex, indicating an important role of higher order areas for goal-directed visual information processing.

## Methods

### Participants

A combined total of 72 participants [49 females, 23 males, mean ± standard-error-of-the-mean (SEM) age = 24.0 ± 0.60 years old; 70 right-handed, 2 left-handed] with normal or corrected-to-normal vision was recruited. Participants gave informed written consent prior to participation. The study was approved by the internal review board of the University of Regensburg. Thirty-six participants were recruited for the main imaging experiment, among which thirty participants completed the GABA measurement and a subset of twenty-four participants also volunteered to complete the Glx measurement. Six additional new participants were recruited for the Glx measurement. Twenty participants completed the control behavioral experiments. Twenty-one participants including five participants from the main imaging experiment were recruited for the control imaging experiments, among which nineteen participants completed both the target enhancement and distractor suppression experiments. One participant was only available for the target enhancement experiment. Another participant was only available for the distractor suppression experiment.

### MOT Task in Main Imaging Experiment

For the main imaging experiment, a MOT task with two attentional load conditions (tracking two targets = low load; tracking four targets = high load) was used (Cavanagh & Alvarez, 2005; Meyerhoff et al., 2017; Pylyshyn & Storm, 1988) (Figure 1). Each trial consisted of cueing, tracking and response phases. The tracking phase was always 12 s-long. The cueing and response phases were each 3 s-long in the GABA measurement and 2 s-long in the Glx measurement. During the cueing phase a total of twelve disks (diameter: 0.5°) was arranged circularly on a black screen (24 x 18° in size). Disks remained stationary during the cueing phase. A subset of disks was highlighted in green as targets while the other disks (i.e., the distractors) were displayed in white. During the tracking phase, the highlighting was turned off and all disks moved independently with a speed of 4.5°/s in different and unpredictable directions across the screen. The motion trajectory of each disk was constrained to avoid collisions with other disks and overlap with the central fixation spot. Disks were repelled from the borders of the screen. For each trial and session trajectories were calculated in advance and randomly drawn (without replacement) from a predefined pool of unique trajectories that was used for each participant. Participants were instructed to track the targets with their attention and maintain central fixation. Each trial terminated with a response phase. During the response phase all disks remained stationary at their final position at the end of the tracking phase and one disk was highlighted in green. Participants indicated whether the highlighted disk was a target or a distractor by pressing one of two buttons on the MRI-safe buttonbox. In half of all trials, the highlighted disk was a target, resulting in a chance level of 50% correct response accuracy. Participants were instructed to respond as quickly and accurately as possible. No feedback about participants’ response accuracy was provided.

**Figure 1.**
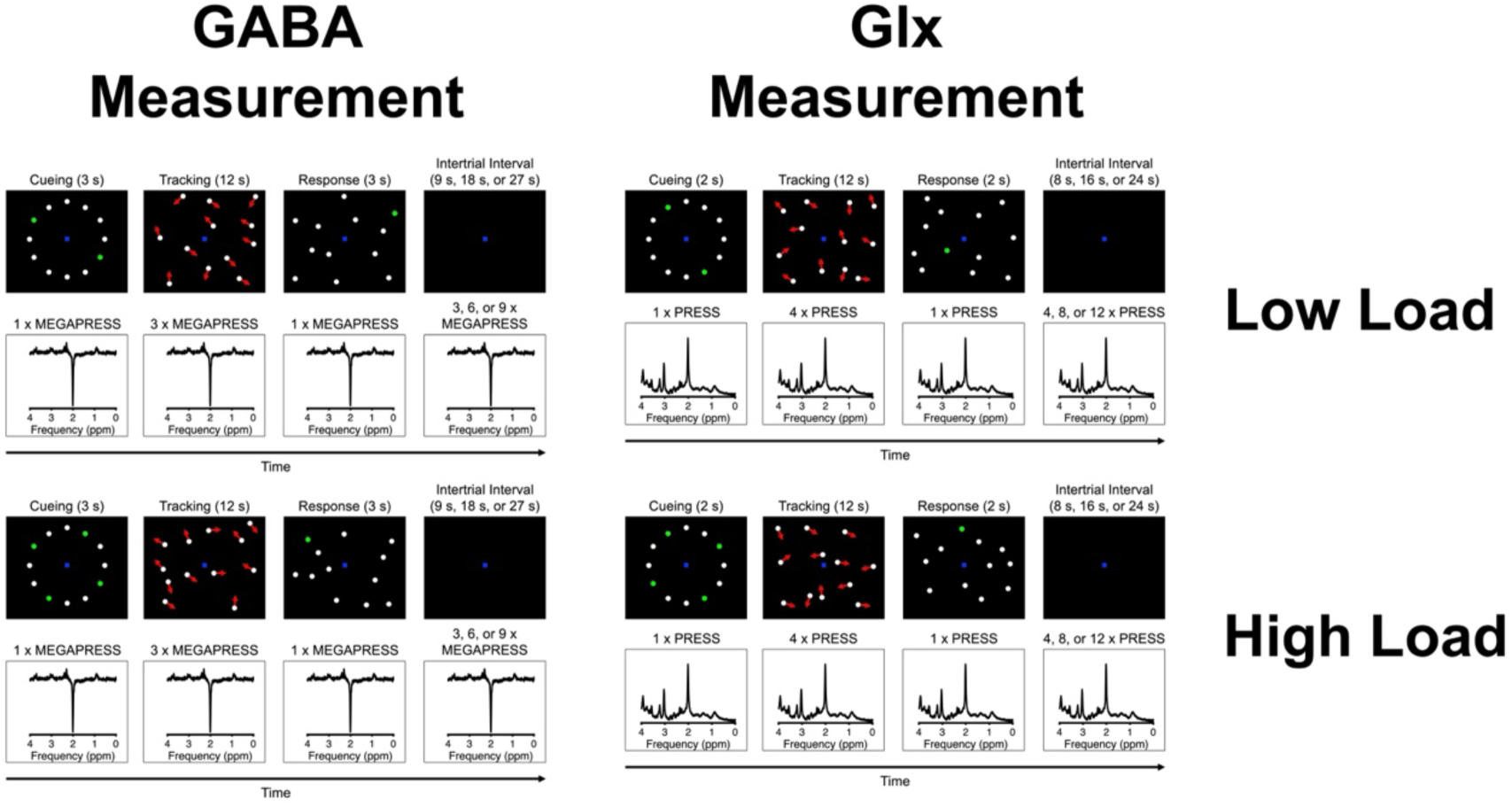
Design of the main imaging experiment. Example trials of the multiple object tracking (MOT) task with low and high attentional load conditions corresponding to tracking two and four targets among distractors, respectively. γ-aminobutyric acid (GABA) was measured with MEGAPRESS scans. Glx (glutamate and its precursor glutamine) was measured with PRESS scans.

**Figure 2.**
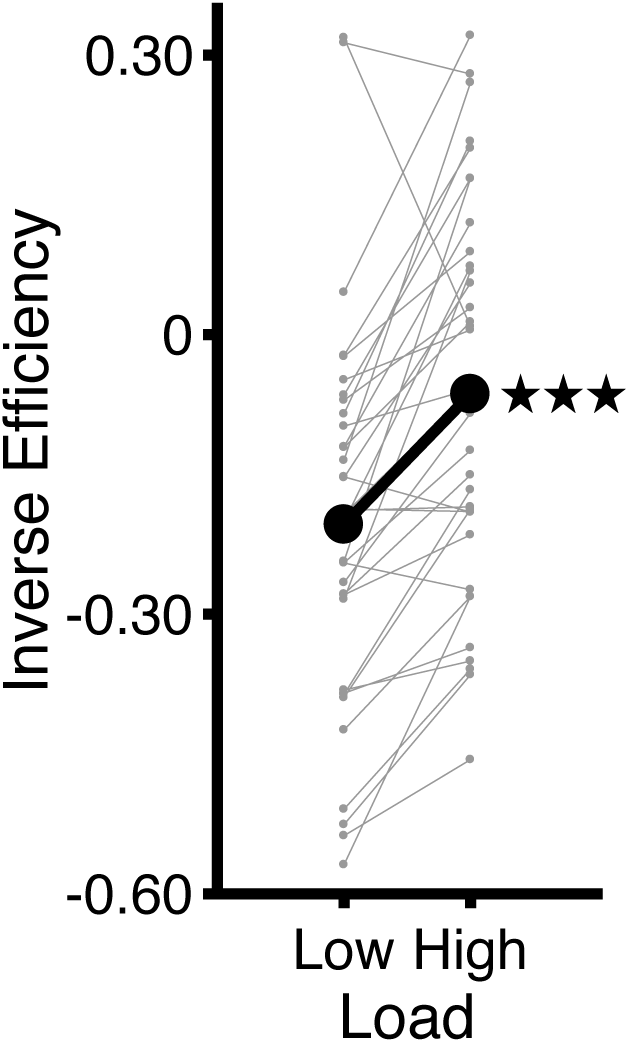
Tracking performance across fMRS runs in the main imaging experiment. The thick line shows mean results across participants. Thin lines show results for each participant. The lower the inverse efficiency score, the faster and more accurate participants responded. The result of a paired-sample *t*-test between tracking with high load and low load is denoted by asterisks. *** *p* < 0.001.

### Main Imaging Experiment

The main imaging experiment included separate fMRS runs for GABA and Glx measurements in the parietal and visual cortex during MOT. A checklist with all fMRS acquisition parameters, analyses and quality checks, as proposed by Choi et al. (2021), is provided in Table 1. In each fMRS run participants performed MOT with low and high load conditions while a time-series of fMRS scans was collected (see Figure 1). Low and high load conditions were presented in random order within each fMRS run. For each load condition there was a total of 18 trials in the GABA measurement and 9 trials in the Glx measurement. For fMRS a mixed design was used in which each trial with MOT was followed by a jittered intertrial interval. This intertrial interval ranged between 9 s, 18 s, and 27 s in the GABA measurement and between 8 s, 16 s, and 24 s in the Glx measurement. During the intertrial interval a central fixation spot on an otherwise blank screen was displayed. Each fMRS run commenced with an on-ramp and terminated with an off-ramp. During ramps a central fixation spot on an otherwise blank screen was displayed. Ramps were 60 s-long in the GABA measurement and 64 s-long in the Glx measurement.

**Table 1.** fMRS checklist proposed by Choi et al. (2021).

|  |  |
| --- | --- |
| **Site (Name or Number):** Brain Imaging Center at University of Regensburg / Bezirksklinikum Regensburg |  |
| **1. Hardware** |  |
| a. Field strength [T] | 3 T |
| b. Manufacturer | Siemens |
| c. Model (software version if available) | Prisma |
| d. RF coils: nuclei (transmit/receive), number of channels, type, body part | 64 channel head/neck coil |
| e. Additional hardware | N/A |
| **2. Acquisition** |  |
| a. Pulse sequence | Siemens MEGAPRESS sequence; Siemens PRESS sequence |
| b. Volume of Interest (VOI) locations | Parietal lobes; occipital lobes |
| c. Nominal VOI size [cm<sup>3</sup>, mm<sup>3</sup>] | 2 x 2 x 4 cm (parietal lobes); 2 x 2 x 2 cm (occipital lobes) |
| d. Repetition Time (TR), Echo Time (TE) [ms, s] | MEGAPRESS (for each of Edit On & Edit Off): TR 1500 ms; TE = 68 ms; PRESS: TR 2000 ms; TE = 30 ms |
| e. Total number of Excitations or acquisitions per spectrum | MEGAPRESS: 472 scans in total, which corresponded to the following experimental conditions (summed across all time-points): on-ramp: 20 scans; cueing phase low load/low enhancement/low suppression: 18 scans; tracking phase low load/low enhancement/low suppression: 72 scans; response phase low load/low enhancement/low suppression: 18 scans; cueing phase high load/high enhancement/high suppression: 18 scans; tracking phase high load/high enhancement/high suppression: 72 scans; response phase high load/high enhancement/high suppression: 18 scans; intertrial interval 216 scans; off-ramp: 20 scans. For each participant spectra for the tracking phases in each condition were averaged across trials with correct participant response.<br>PRESS: 352 scans in total, which corresponded to the following experimental conditions (summed across all time-points): on-ramp: 32 scans; cueing phase low load/low enhancement/low suppression: 9 scans; tracking phase low load/low enhancement/low suppression: 54 scans; response phase low load/low enhancement/low suppression: 9 scans; |
|  | cueing phase high load/high enhancement/high suppression: 9 scans; tracking phase high load/high enhancement/high suppression: 54 scans; response phase high load/high enhancement/high suppression: 9 scans; intertrial interval 144 scans; off-ramp: 32 scans. For each participant spectra for the tracking phases in each condition were averaged across trials with correct participant response. |
| f. Additional sequence parameters (spectral width in Hz, number of spectral points, frequency offsets) | Spectral bandwidth: 1200 Hz; Spectral points: 1024 |
| g. Water Suppression Method | WET |
| h. Shimming Method, reference peak, and thresholds for “acceptance of shim” chosen | Automated B<sub>0</sub> field mapping to < 20 Hz |
| i. Triggering or motion correction method | N/A |
| **3. Data analysis methods and outputs** |  |
| a. Analysis software | LC Model Version 6.3-R; Osprey Version 2.4.0 |
| b. Processing steps deviating from quoted reference or product | N/A |
| c. Output measure | Tissue-corrected concentrations relative to NAA from the same spectrum |
| d. Quantification references and assumptions, fitting model assumptions | Basis sets were imported from LC model software package and included the following spectra: MEGAPRESS difference: γ-Aminobutyric Acid (GABA), Glutamine (Gln), Glutamate (Glu), Glutathione (GSH), N-Acetylaspartate (NAA), N-Acetylaspartylglutamate (NAAG). PRESS: Aspartate (Asp), Creatine (Cr), Phosphocreatine (PCr), γ-Aminobutyric Acid (GABA), Glucose (Glc), Glutamine (Gln), Glutamate (Glu), Glycerophosphocholine (GPC), Glutathione (GSH), myo-Inositol (Ins), L-Lactate (Lac), N-Acetylaspartate (NAA), N-Acetylaspartylglutamate (NAAG), scyllo-Inositol (Scyllo), Taurine (Tau). |
| **4. Data Quality** |  |
| a. Reported variables | CRLB, shimming, S/N |
| b. Data exclusion criteria | CRLB > 25%, shimming ≥ 20 Hz, S/N < 5 |
| c. Quality measures of postprocessing Model fitting | CRLB, S/N |
| d. Sample Spectrum | See Supplementary Figures 1-4 |

### MOT Tasks in Control Experiments

For the control behavioral and imaging experiments, two modified versions of the MOT task were used, which were based on previously proposed designs to separately manipulate levels of target enhancement and distractor suppression during tracking (Bettencourt & Somers, 2009) (Figure 5). Using these modified designs, Bettencourt and Somers (2009) found that higher levels of target enhancement and distractor suppression each led to a decrease in tracking performance. Moreover, they showed that this change in behavioral performance did not result from stronger crowding during tracking with high than low target enhancement and distractor suppression (Bettencourt & Somers, 2009). All parameters in the modified versions of the MOT task including motion parameters, trial duration and number of trials with target vs. distractor highlighted in the response phase were identical with the MOT task in the main imaging experiment except for the following aspects: For the target enhancement experiment, a MOT task with two different sizes of all disks was used (large disk size = low enhancement; small disk size = high enhancement). A total of eight disks were presented in each trial. Four disks were highlighted as targets during the cueing phase. The remaining four disks were distractors. For the low and high enhancement conditions, the disk sizes were varied between 1.25° and 0.25°, respectively. For the distractor suppression experiment, a MOT task with two different numbers of distractor disks was used (low number of distractors = low suppression; high number of distractors = high suppression). For the low suppression condition, four out of eight disks were highlighted as targets during the cueing phase yielding a total of four distractors. For the high suppression condition, four out of twelve disks were highlighted as targets yielding a total of eight distractors.

### Control Behavioral Experiments

Two control behavioral experiments were carried out to establish behavioral measurements of target enhancement and distractor suppression during MOT as in Bettencourt & Somers (2009). The results were also used to determine the number of participants required for the control imaging experiments. The control behavioral experiments were conducted in a single session in a psychophysics-dedicated room with room lights turned off. Participants used a chin rest and responded by pressing one of two keys on the keyboard to indicate whether the highlighted disk in the response phase was a target or a distractor. Participants completed two runs for each of the target enhancement and distractor suppression experiments using the same number of trials as in the GABA and Glx measurements in the main imaging experiment. For the control behavioral experiments, the intertrial interval was shortened to 2 s and on-/off-ramps were omitted.

### Control Imaging Experiments

Two control imaging experiments were carried out to measure GABA and Glx concentrations in parietal cortex during MOT with low and high levels of target enhancement and distractor suppression, respectively. The MOT designs were identical with the control behavioral experiments. The number of trials, intertrial intervals and on-/off-ramps for GABA and Glx measurements were identical with the main imaging experiment. The results of the control behavioral experiments were used to calculate the number of participants necessary to achieve a power of 0.95 at alpha = 0.001 (two-tailed) using a paired-sample *t*-test to detect a minimum effect size of interest for a difference in inverse efficiency score between high and low levels of target enhancement and distractor suppression, respectively. The results of the power analysis (using G*Power; Faul et al., 2007) showed that a total of fifteen participants was necessary to detect a minimum effect size of interest (*d* = 1.65, calculated from the control behavioral experiment) for a difference between high and low target enhancement. A total of thirteen participants was necessary to detect a minimum effect size of interest (*d* = 1.85, calculated from the control behavioral experiment) for a difference between high and low distractor suppression. Based on these results, we decided to run the same number of participants in the control imaging experiments as in the control behavioral experiments (i.e., twenty participants).

### Scanning Parameters

In the main imaging experiment, single-voxel proton (^1^H) fMRS was conducted for two types of fMRS scans (MEGAPRESS for GABA measurement and PRESS for Glx measurement) and volumes-of-interest (VOIs; located in the parietal and occipital lobes, respectively). In the control imaging experiments, MEGAPRESS and PRESS scans were collected for the parietal VOI.

MEGAPRESS scans (Mescher et al., 1996, 1998) were acquired with alternating Edit On and Edit Off scans with the following parameters: time-to-repeat (TR) = 1.5 s; time-to-echo (TE) = 68 ms; flip angle (FA) = 90°. The total number of scans (Edit On and Edit Off combined) for each MEGAPRESS fMRS run was 472. A frequency selective, single band Gauss pulse was utilized to saturate the *β*-CH_2_ signal at 1.94 ppm and to refocus the J evolution of the triplet γ-CH_2_ resonance of GABA at 3 ppm (‘Edit On’). The very same Gauss pulse was used to irradiate the opposite part of the spectrum at 7.46 ppm (‘Edit Off’). The ‘Edit Off’ spectrum was subtracted from the ‘Edit On’ spectrum to produce a difference spectrum. For each load condition in the main imaging experiment and for each level of target enhancement and distractor suppression in the control imaging experiments, there were a total of 18 scans for the cueing phase, 72 scans for the tracking phase, and 18 scans for the response phase. The remaining scans corresponded to the intertrial interval (216 scans) and on-/off-ramps (20 scans for each ramp). The total number of scans collected during the tracking phase in each condition was based on previous studies (Floyer-Lea et al., 2006; Frank et al., 2022), which used 64 scans to estimate GABA from a MEGAPRESS difference spectrum.

PRESS scans (Bottomley et al., 1984; Bottomley & Hardy, 1987) were acquired with the following parameters: TR = 2 s; TE = 30 ms; FA = 90°. The total number of scans for each PRESS fMRS run was 352. For each load condition in the main imaging experiment and for each level of target enhancement and distractor suppression in the control imaging experiments, there were a total of 9 scans for the cueing phase, 54 scans for the tracking phase, and 9 scans for the response phase. The remaining scans corresponded to the intertrial interval (144 scans) and on-/off-ramps (32 scans for each ramp).

WET water suppression was used (Ogg et al., 1994). In the main imaging experiment, a total of four fMRS runs were conducted corresponding to MEGAPRESS for the parietal VOI, MEGAPRESS for the visual VOI, PRESS for the parietal VOI, and PRESS for the visual VOI. In the control imaging experiments, a total of four fMRS runs were conducted for the parietal VOI corresponding to MEGAPRESS for target enhancement, MEGAPRESS for distractor suppression, PRESS for target enhancement, and PRESS for distractor suppression. The parietal VOI had a size of 2 x 2 x 4 cm and was placed approximately perpendicular to the intraparietal sulcus. The visual VOI had a size of 2 x 2 x 2 cm and was placed perpendicular to the calcarine sulcus. Proximity to the dura was avoided during VOI placement to minimize macromolecule contamination. See Figure 3 for VOI locations in a representative participant. VOIs were centered between the two hemispheres. Non-neural tissue containing lipids was avoided during VOI placement. For the purpose of VOI placement, three high-resolution anatomical scans were acquired for the coronal, sagittal and transverse planes prior to fMRS. These anatomical scans had the following parameters: coronal plane: TR = 0.25 s, TE = 2.46 ms, FA = 70°, acquisition matrix (AM) = 288 x 288, 35 slices, voxel size = 0.8 x 0.8 x 4.0 mm, inter-slice gap = 1.20 mm; sagittal plane: TR = 0.19 s, TE = 2.46 ms, FA = 70°, AM = 288 x 288, 25 slices, voxel size = 0.8 x 0.8 x 4.0 mm, inter-slice gap = 1.20 mm; transverse plane: TR = 0.19 s, TE = 2.46 ms, FA = 70°, AM = 288 x 288, 27 slices, voxel size = 0.8 x 0.8 x 4.0 mm, inter-slice gap = 1.20 mm. After each fMRS run, a water reference scan without water suppression was acquired for the same VOI using the following parameters: MEGAPRESS: TR = 1.5 s; TE = 30 ms; FA = 90°; 8 scans; PRESS: TR = 2 s; TE = 30 ms; FA = 90°; 16 scans.

**Figure 3.**
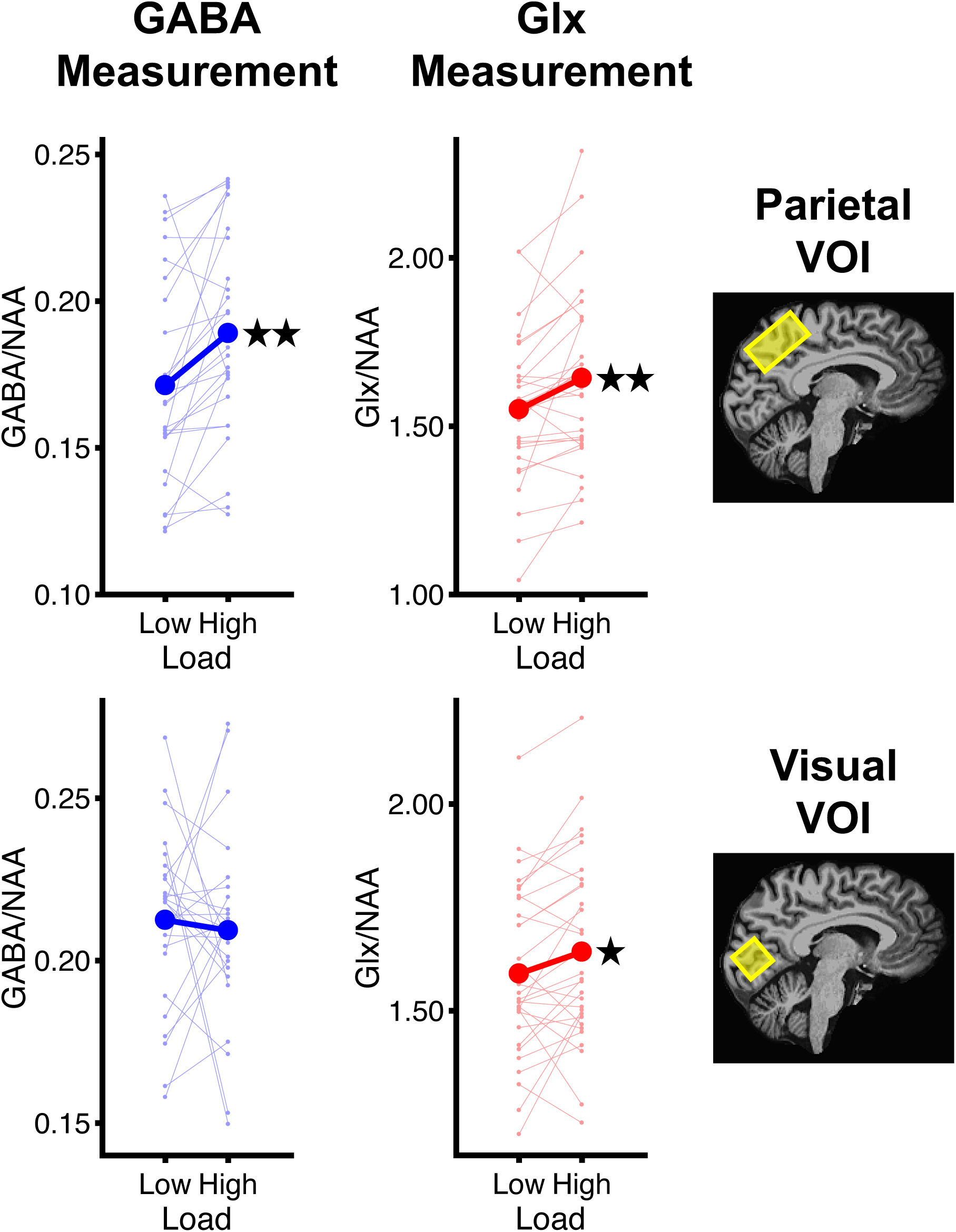
Metabolite concentrations in the main imaging experiment. Thick lines show mean results across participants for the parietal and visual volumes-of-interest (VOIs). Thin lines show results for each participant. GABA and Glx concentrations are normalized to the concentration of N-acetyl-aspartate (NAA). Results of paired-sample *t*-tests between tracking with high load and low load are denoted by asterisks. * *p* < 0.05, ** *p* < 0.01.

Automatic shimming (gradient field adjustments to increase the homogeneity of the magnet field B_0_) was conducted prior to each fMRS run. For each fMRS run and participant the shim values were kept below a full-width at half-maximum of 20 Hz. In the main imaging experiment, the mean ± SEM shim values across participants were as follows: MEGAPRESS: 13.6 ± 0.26 Hz (parietal VOI) and 13.8 ± 0.18 Hz (visual VOI); PRESS: 13.6 ± 0.22 Hz (parietal VOI) and 14.5 ± 0.26 Hz (visual VOI); water reference scan in MEGAPRESS: 13.9 ± 0.31 Hz (parietal VOI) and 13.9 ± 0.21 Hz (visual VOI); water reference scan in PRESS: 13.7 ± 0.28 Hz (parietal VOI) and 14.5 ± 0.24 Hz (visual VOI). In the control imaging experiments, the values for the parietal VOI were as follows: target enhancement: MEGAPRESS: 14.1 ± 0.39 Hz; PRESS: 14.0 ± 0.32 Hz; water reference scan in MEGAPRESS: 14.9 ± 0.55 Hz; water reference scan in PRESS: 14.2 ± 0.34 Hz; distractor suppression: MEGAPRESS: 15.2 ± 0.46 Hz; PRESS: 14.8 ± 0.47 Hz; water reference scan in MEGAPRESS: 15.0 ± 0.42 Hz; water reference scan in PRESS: 14.8 ± 0.45 Hz.

In order to calculate volume fractions for each VOI (see *Imaging Analysis* below) a high-resolution anatomical scan of the brain was collected using a magnetization prepared rapid gradient echo sequence with the following scanning parameters: TR= 2.3 s, TE = 2.32 ms, FA = 8°, AM = 256 x 256, 192 sagittal slices, voxel size = 0.9 x 0.9 x 0.9 mm, interslice gap = 0.45 mm. For a subset of five participants, this scan could not be collected due to time constraints and the sagittal scan acquired for VOI placement was used instead.

Some studies reported that the concentration of GABA is modulated by the menstrual cycle (De Bondt et al., 2015; Duncan et al., 2014). Therefore, in female participants not using contraception, the experiments were only conducted during the follicular phase.

### Stimulus Presentation

Stimuli were generated using Psychtoolbox (Brainard, 1997; Pelli, 1997) running in Matlab (The Mathworks, Natick, MA, USA). In the scanner, stimuli were presented using a projector with gamma-correction onto a translucent screen located at the back of the scanner bore. Participants viewed the screen with a headcoil-mounted mirror. Outside the scanner, stimuli were presented on an LCD-screen with gamma-correction.

### Behavioral Analysis

Response accuracy and median response time (in seconds) were combined into the inverse efficiency score for each run and participant (Townsend & Ashby, 1978, 1983). To this purpose median response time was divided by response accuracy (in proportion of one) and the result was log-transformed (Bruyer & Brysbaert, 2011). The inverse efficiency score was then combined across runs by averaging for each participant in each experiment. For correlational analyses between fMRS and behavioral results, the inverse efficiency score of the corresponding fMRS run was used. For one participant in the main imaging experiment, response accuracy was at chance level in the GABA measurement. The behavioral and GABA results from this participant were excluded from all further analyses. For one participant in the control imaging experiments, response accuracy was at chance level in all fMRS runs. The behavioral and fMRS results from this participant were excluded from all further analyses.

### Imaging Analysis

The fMRS data were analyzed using the LC Model (Provencher, 1993, 2001). Analysis steps as in previous studies were applied (Frank et al., 2023). Details are reported in Table 1 (Choi et al., 2021; Lin et al., 2021). Before running the analysis, spectra from different scans in the fMRS time-series were averaged across trials and conditions as follows. Average spectra were produced for the tracking phase in each condition for trials with correct participant response (Supplementary Figures 1-4). The mean ± SEM number of spectra used for averaging across participants in the main imaging experiment was as follows: MEGAPRESS: 69.7 ± 0.81 (parietal VOI) and 70.5 ± 0.46 (visual VOI) in low load; 63.3 ± 1.45 (parietal VOI) and 61.7 ± 1.58 (visual VOI) in high load; PRESS: 50.0 ± 1.01 (parietal VOI) and 51.4 ± 0.80 (visual VOI) in low load; 47.8 ± 1.33 (parietal VOI) and 48.2 ± 0.97 (visual VOI) in high load. The number of spectra used in the control imaging experiments was as follows: MEGAPRESS: 68.8 ± 1.25 (low enhancement) and 65.7 ± 2.08 (high enhancement); 65.5 ± 1.65 (low suppression) and 61.9 ± 2.23 (high suppression); PRESS: 50.5 ± 1.24 (low enhancement) and 47.1 ± 1.40 (high enhancement); 49.9 ± 1.66 (low suppression) and 47.7 ± 1.34 (high suppression).

The averaged MEGAPRESS difference spectra were analyzed in the chemical shift range between 1.95 and 4.2 ppm. The averaged PRESS spectra were analyzed in the chemical shift range between 0.2 and 4.0 ppm. Water scaling and eddy current correction were performed. The metabolite intensities were fitted to a linear combination of spectra of individual metabolites derived from imported metabolite basis sets (Provencher, 1993, 2001; Table 1).

The following metabolite concentrations estimated for the tracking phase in each condition and fMRS run were further analyzed: MEGAPRESS: GABA and N-acetyl-aspartate (NAA). NAA is a marker of neuronal density and mitochondrial function (Schuff et al., 2006) and is commonly used as a control metabolite for normalization due to its stability over experimental conditions and scan time (Duncan et al., 2014; Frank et al., 2023; Mullins et al., 2014; Stagg, 2014). PRESS: Glx and NAA. Prior to normalization, metabolite concentrations were corrected for each participant’s volume fractions inside the fMRS VOIs. Volume fractions within the fMRS VOIs corresponding to gray matter, white matter and other types of volume including cerebrospinal fluid were calculated using the Freesurfer segmentation of each participant’s reconstructed high-resolution anatomical scan of the brain (Dale et al., 1999; Fischl et al., 1999). Across participants and experiments the mean ± SEM volume fractions were as follows: Gray matter: 61.7 ± 0.96% (parietal VOI) and 60.1 ± 0.90% (visual VOI); white matter: 15.9 ± 0.65% (parietal VOI) and 34.1 ± 0.89% (visual VOI); other types of volume: 22.4 ± 1.17% (parietal VOI) and 5.85 ± 0.59% (visual VOI). Volume fraction correction was performed using the approach proposed Kolasinski et al. (2017). The concentrations of GABA and Glx were corrected for the proportion of gray matter within each VOI by dividing by [gray matter / (gray matter + white matter + other)]. The concentration of NAA was corrected for the proportion of total brain volume within each VOI by dividing by [(gray matter + white matter) / (gray matter + white matter + other)]. Thereafter, the corrected concentration of GABA was normalized by dividing it by the corrected concentration of NAA extracted from the same MEGAPRESS difference spectrum. The corrected concentration of Glx was normalized by dividing it by the corrected concentration of NAA extracted from the same PRESS spectrum.

Post-hoc analyses were performed to calculate which brain regions were included in the gray matter of each VOI. The PALS segmentation (Van Essen, 2005) was remapped from the Freesurfer standard brain to each participant’s individual brain and the percentage of Brodmann areas in the gray matter of each VOI was calculated for each participant. Across participants and experiments, approximately seventy percent of gray matter within the parietal VOI corresponded to Brodmann areas 1, 2, 3, 5, 7 and 31 (mean ± SEM: areas 1-3: 7.33 ± 0.69%; area 5: 22.1 ± 1.08%; area 7: 20.2 ± 1.25%; area 31: 16.9 ± 1.31%). The remaining gray matter corresponded mostly to the medial part of Brodmann area 4 (32.0 ± 2.39%). Within the gray matter of the visual VOI, 60.4 ± 2.10% corresponded to Brodmann area 17 and 37.5 ± 1.95% corresponded to Brodmann area 18 across participants. Gray matter corresponding to other areas was negligible. The mean ± SEM MNI coordinates of the parietal and visual VOIs across participants and experiments were as follows: Parietal VOI: X = 0.04 ± 0.23, Y = –47.9 ± 0.76, Z = 52.7 ± 0.54; Visual VOI: X = 0.39 ± 0.31, Y = –81.4 ± 0.42, Z = 4.39 ± 0.54.

We only included spectral results for which the signal-to-noise (SN) ratio as estimated by the LC model was higher than 5% (Wilson et al., 2019). None of a participant’s spectra had to be excluded. The mean SN ratios (± SEM) across participants’ spectra in the main imaging experiment were as follows: MEGAPRESS: 24.8 ± 1.42% (parietal VOI) and 16.2 ± 0.54% (visual VOI) in low load; 23.1 ± 1.14% (parietal VOI) and 15.2 ± 0.51% (visual VOI) in high load; PRESS: 30.2 ± 1.36% (parietal VOI) and 28.7 ± 0.83% (visual VOI) in low load; 29.7 ± 1.39% (parietal VOI) and 28.1 ± 0.85% (visual VOI) in high load. The values in the control imaging experiments for the parietal VOI were as follows: MEGAPRESS: 22.3 ± 1.84% (low enhancement) and 21.5 ± 1.91% (high enhancement); 24.2 ± 2.01% (low suppression) and 24.2 ± 1.97% (high suppression); PRESS: 30.5 ± 2.30% (low enhancement) and 30.4 ± 2.24% (high enhancement); 29.9 ± 2.23% (low suppression) and 29.4 ± 2.17% (high suppression).

The Cramer-Rao Lower Bounds (CRLB) indicate the reliability of quantification of GABA, Glx, and NAA. A criterion of 25% was used to reject low-quality results (following the approach in Frank et al., 2022). In the main imaging experiment, GABA results for the parietal VOI in three participants and for the visual VOI in four participants exceeded the inclusion criterion. GABA and NAA results from these participants in MEGAPRESS were excluded. Therefore, out of twenty-nine participants in MEGAPRESS, GABA and NAA results in the parietal VOI for twenty-six participants and for twenty-five participants in the visual VOI were further analyzed. None of the Glx and NAA results in any VOI in PRESS exceeded the CRLB inclusion criterion. The mean CRLB (± SEM) across included participants in the main imaging experiment were as follows: GABA (MEGAPRESS): 16.8 ± 0.58% (parietal VOI) and 19.4 ± 0.57% (visual VOI) in low load; 15.4 ± 0.42% (parietal VOI) and 20.1 ± 0.49% (visual VOI) in high load; NAA (MEGAPRESS): 3.50 ± 0.16% (parietal VOI) and 3.40 ± 0.10% (visual VOI) in low load; 3.69 ± 0.15% (parietal VOI) and 3.72 ± 0.12% (visual VOI) in high load; Glx (PRESS): 6.47 ± 0.15% (parietal VOI) and 6.73 ± 0.17% (visual VOI) in low load; 6.10 ± 0.14% (parietal VOI) and 6.70 ± 0.13% (visual VOI) in high load; NAA (PRESS): 2.60 ± 0.10% (parietal VOI) and 2.57 ± 0.11% (visual VOI) in low load; 2.67 ± 0.11% (parietal VOI) and 2.70 ± 0.11% (visual VOI) in high load.

In the control imaging experiments for the parietal VOI, GABA results from one participant exceeded the CRLB inclusion criterion. GABA and NAA results from this participant in MEGAPRESS were excluded. Therefore, out of nineteen participants in MEGAPRESS, GABA and NAA results for eighteen participants in MEGAPRESS for target enhancement and distractor suppression were further analyzed. None of the Glx and NAA results in PRESS exceeded the CRLB inclusion criterion. The mean CRLB (± SEM) across included participants in the control imaging experiments were as follows: GABA (MEGAPRESS): 16.1 ± 0.80% (low enhancement) and 16.7 ± 0.87% (high enhancement); 17.1 ± 0.71% (low suppression) and 18.3 ± 0.75% (high suppression); NAA (MEGAPRESS): 4.58 ± 0.59% (low enhancement) and 4.84 ± 0.67% (high enhancement); 3.63 ± 0.29% (low suppression) and 3.52 ± 0.22% (high suppression); Glx (PRESS): 6.68 ± 0.29% (low enhancement) and 6.68 ± 0.32% (high enhancement); 6.84 ± 0.36% (low suppression) and 6.84 ± 0.41% (high suppression); NAA (PRESS): 2.94 ± 0.18% (low enhancement) and 2.94 ± 0.16% (high enhancement); 3.05 ± 0.34% (low suppression) and 3.05 ± 0.25% (high suppression).

### Statistics

The results were analyzed using *t*-tests and Pearson correlations. The two-tailed alpha-level was set to 0.05. Cohen’s *d* and Pearson’s *r* were calculated as measures of effect size for *t*-test and Pearson correlation, respectively. Cohen’s *d* for repeated measures was calculated using the formula implemented in G*Power (Faul et al., 2007).

## Results

### Main Imaging Experiment

The behavioral results showed that the inverse efficiency score was significantly greater in MOT with high load than with low load [*t*(34) = 5.79, *p* < 0.001, *d* = 0.98], suggesting that participants responded slower and were less accurate with increasing tracking load (Figure 2 and Supplementary Figure 5). The fMRS results showed that the concentration of GABA was significantly greater in MOT with high load than with low load in the parietal VOI [*t*(25) = 3.25, *p* = 0.003, *d* = 0.64] but not in the visual VOI [*t*(24) = −0.84, *p* = 0.41] (Figure 3). The concentration of Glx was significantly greater in MOT with high load than with low load in both VOIs [parietal VOI: *t*(29) = 3.27, *p* = 0.003, *d* = 0.60; visual VOI: *t*(29) = 2.37, *p* = 0.02, *d* = 0.43] (Figure 3).

The fMRS and behavioral results in the parietal VOI were correlated (Figure 4): The greater was the increase in GABA concentration from low to high load, the smaller was the difference in inverse efficiency score between load conditions (i.e., the smaller was the decrease in response accuracy and increase in response time with increasing tracking load) across participants (*r* = −0.47, *p* = 0.02). A different association was found between Glx concentration in the parietal VOI and tracking performance (Figure 4): The greater the increase in Glx concentration from low to high load, the greater was the increase in inverse efficiency score with load (i.e., the worse was response accuracy and the slower was response time with increasing tracking load) across participants (*r* = 0.49, *p* = 0.006). There was no significant correlation with tracking performance in the visual VOI (GABA: *r* = −0.17, *p* = 0.43; Glx: *r* = 0.24, *p* = 0.21) (Figure 4).

**Figure 4.**
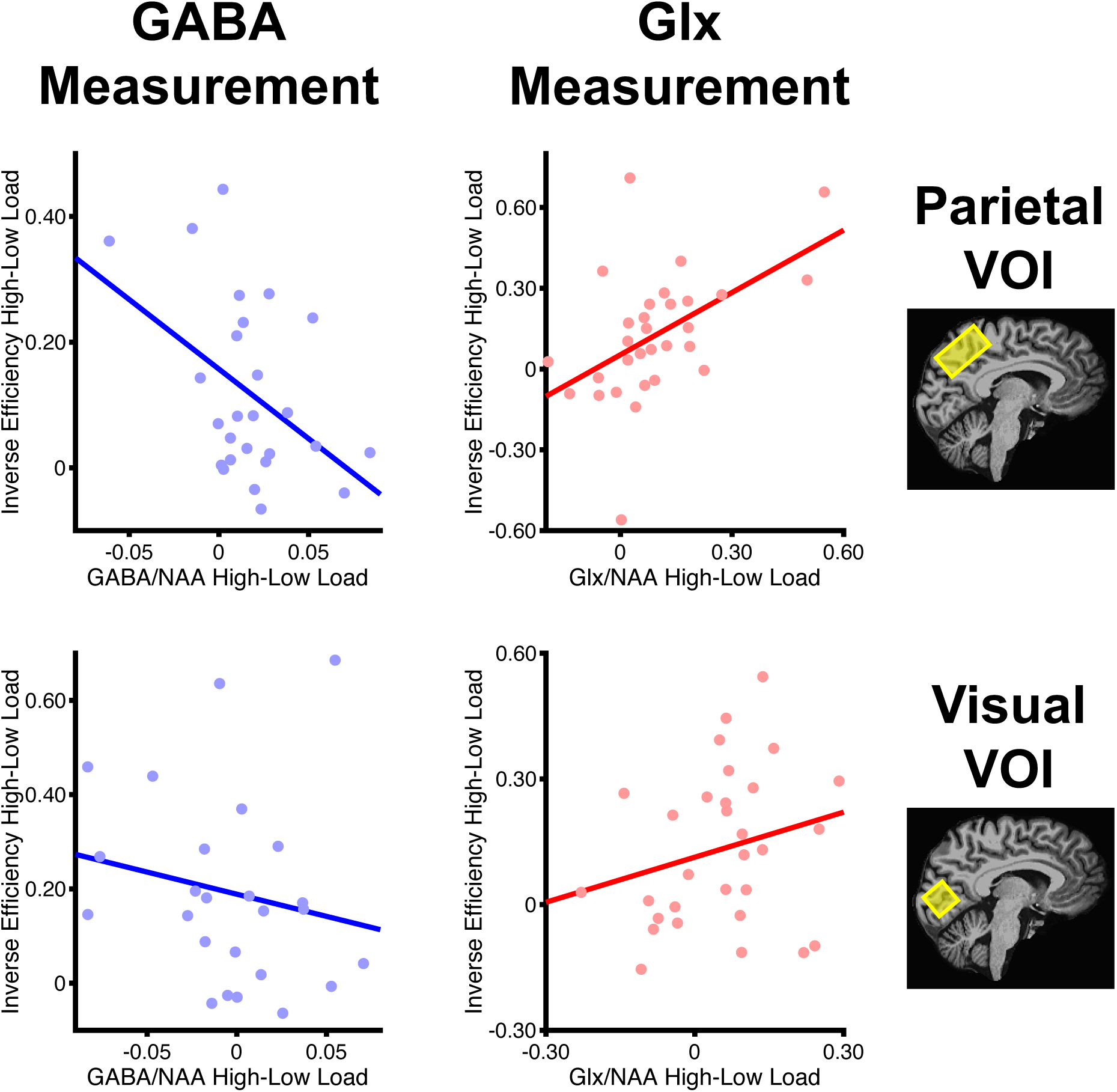
Correlational results in the main imaging experiment. Load-dependent changes in GABA and Glx concentrations during tracking were calculated by subtracting low load from high load concentrations for each participant. More positive values on the x-axis correspond to greater concentration increase from low to high load. Load-dependent changes in tracking performance were calculated by subtracting the inverse efficiency score in low load from high load for each participant. More positive values on the y-axis correspond to greater increase in inverse efficiency score from low to high load (i.e., greater decrease of response accuracy and greater increase of response time from low to high load). Each correlational analysis was carried out between metabolite concentrations and tracking performance from the same fMRS run. Each dot shows the result from a different participant.

### Control Behavioral Experiments

The inverse efficiency score was significantly greater in MOT with high than with low target enhancement [*t*(19) = 7.36, *p* < 0.001, *d* = 1.65], suggesting that participants responded less accurately and more slowly when the disk size was small during tracking (Figure 6A and Supplementary Figure 6A,B). Similarly, the inverse efficiency score was significantly greater in MOT with high than with low distractor suppression [*t*(19) = 8.29, *p* < 0.001, *d* = 1.85], suggesting that participants were less accurate and responded more slowly when the number of distractors was high during tracking (Figure 6B and Supplementary Figure 6C,D).

**Figure 5.**
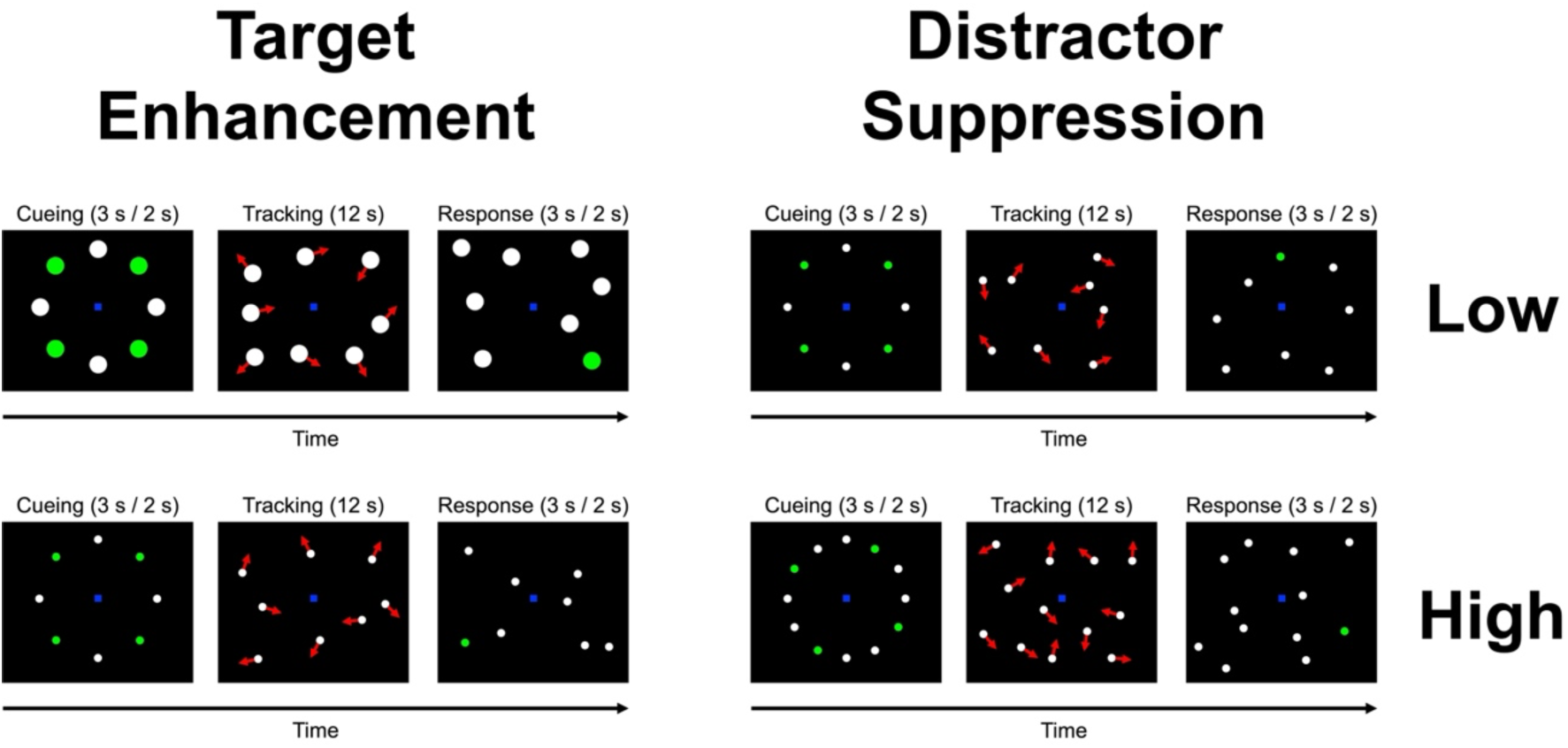
Design of control behavioral and imaging experiments for the measurement of target enhancement and distractor suppression. In the control imaging experiments, GABA and Glx concentrations during MOT were measured in the parietal VOI exactly as in the main imaging experiment with MEGAPRESS and PRESS scans, respectively (see Figure 1). Cueing and response periods were 3 s-long each for MEGAPRESS scans and 2 s-long each for PRESS scans. A jittered intertrial interval was included after each MOT trial exactly as in the main imaging experiment.

**Figure 6.**
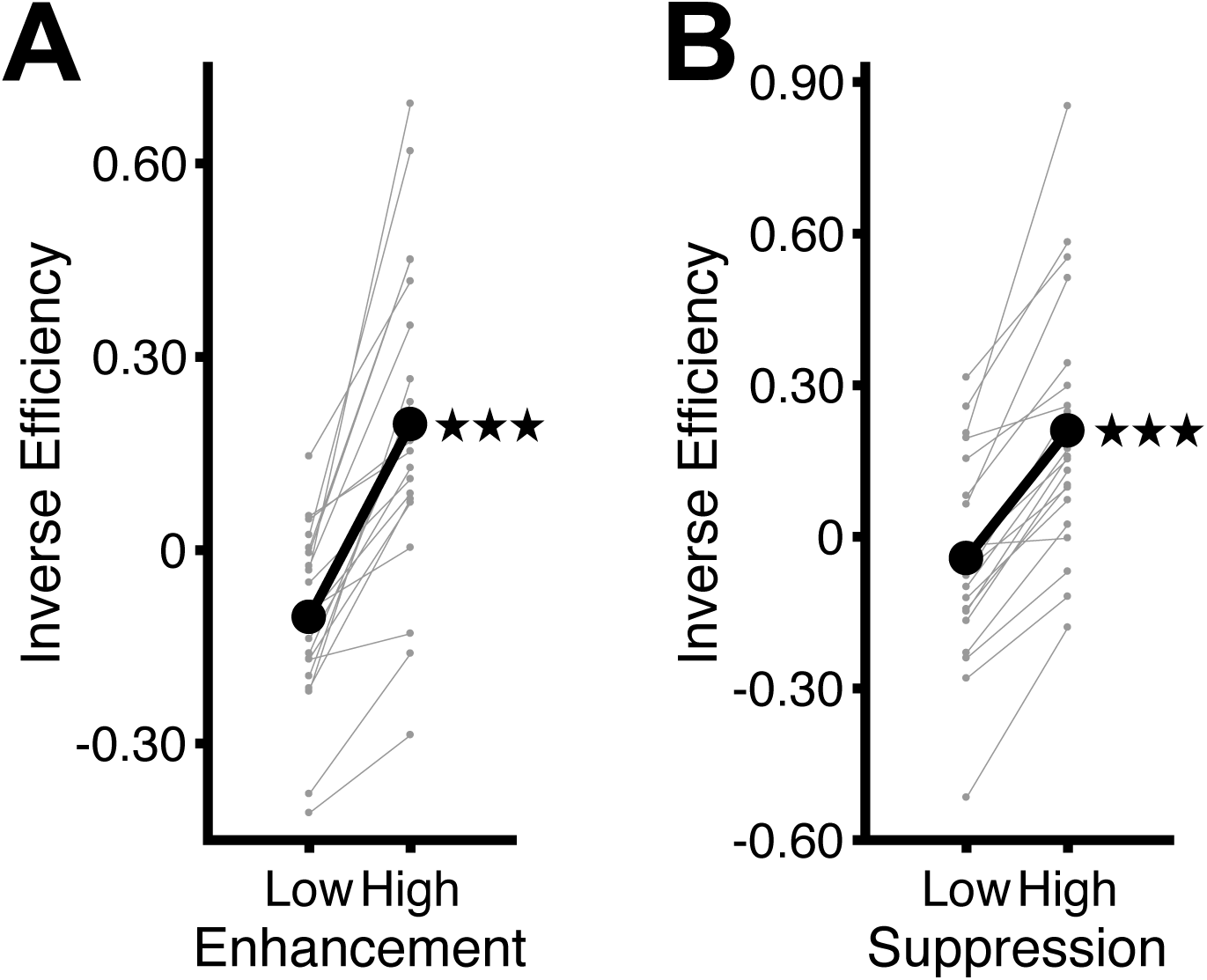
Tracking performance across runs in each control behavioral experiment. (**A**) Target enhancement experiment. (**B**) Distractor suppression experiment. Otherwise same as Figure 2. *** *p* < 0.001.

### Control Imaging Experiments

Results for tracking performance in the control imaging experiments replicated results in the control behavioral experiments as follows: The inverse efficiency score was significantly greater in MOT with high than with low enhancement [*t*(18) = 6.52, *p* < 0.001, *d* = 1.50] and in MOT with high than with low suppression [*t*(18) = 3.61, *p* = 0.002, *d* = 0.83] (Figure 7 and Supplementary Figure 7). The fMRS results showed no significant differences in metabolite concentrations for the between-condition comparison in the target enhancement [GABA: *t*(17) = 0.58, *p* = 0.57; Glx: *t*(18) = −0.69, *p* = 0.50] and distractor suppression [GABA: *t*(17) = −1.50, *p* = 0.15; Glx: *t*(18) = −0.14, *p* = 0.89] experiments (Figure 8). However, the fMRS and behavioral results were correlated (Figure 9). In the target enhancement experiment, the greater was the increase in Glx concentration from low to high enhancement, the smaller was the difference in inverse efficiency score between the two target enhancement conditions (i.e., the smaller was the decrease in response accuracy and increase in response time from low to high enhancement) across participants (*r* = −0.49, *p* = 0.03) (Figure 9). There was no significant correlation between changes in GABA concentration and inverse efficiency score from low to high enhancement (*r* = −0.15, *p* = 0.56) (Figure 9). In the distractor suppression experiment, metabolite concentrations and tracking performance were correlated in similar directions as in the main imaging experiment: The greater was the increase in GABA concentration from low to high suppression, the smaller was the difference in inverse efficiency score between the two distractor suppression conditions across participants (*r* = −0.52, *p* = 0.03) (Figure 9). Glx concentration and tracking performance was correlated in the opposite direction: The greater was the increase in Glx concentration from low to high suppression, the larger was the difference in inverse efficiency score between the two distractor suppression conditions across participants (*r* = 0.46, *p* = 0.049) (Figure 9).

**Figure 7.**
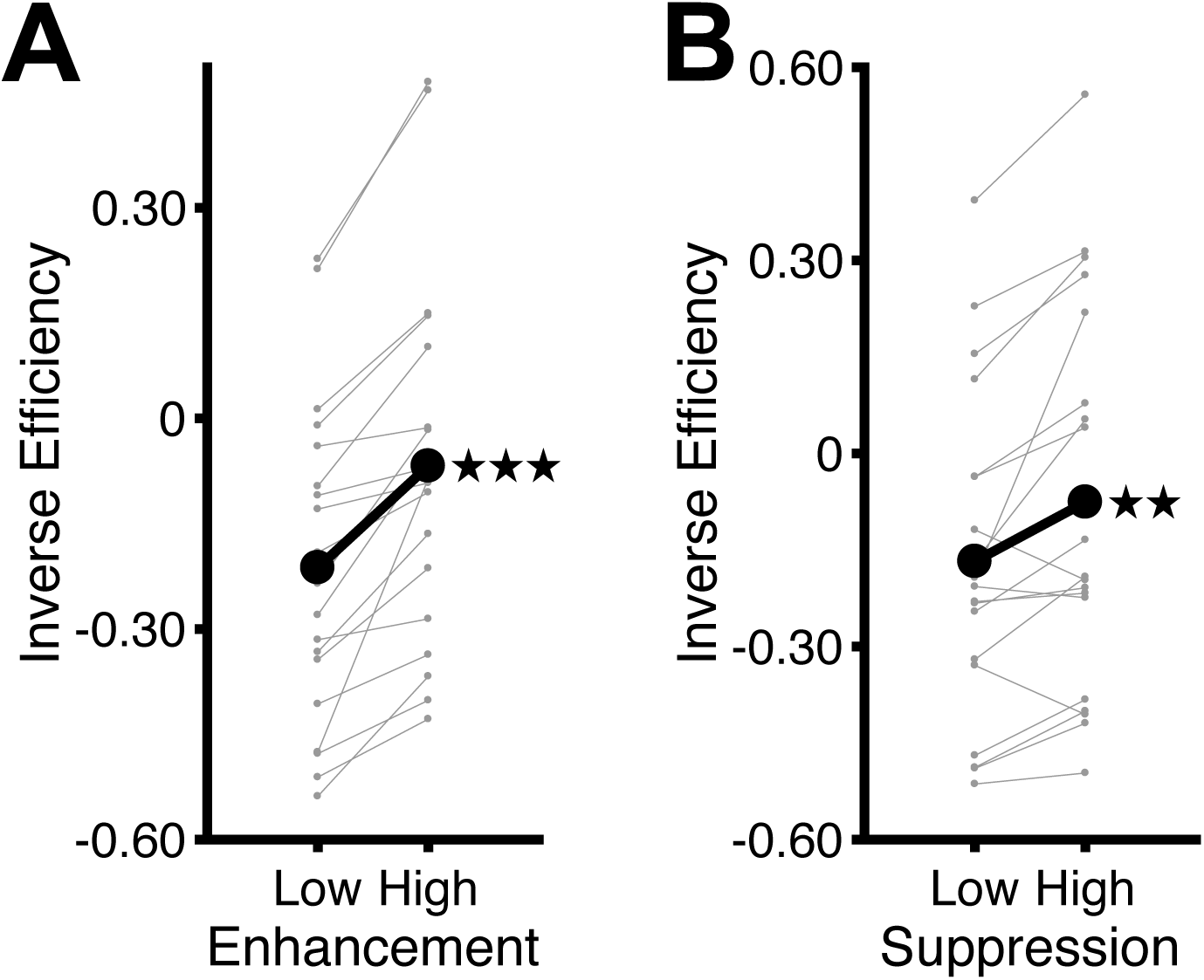
Tracking performance across fMRS runs in each control imaging experiment. (**A**) Target enhancement experiment. (**B**) Distractor suppression experiment. Otherwise same as Figure 6. ** *p* < 0.01, *** *p* < 0.001.

**Figure 8.**
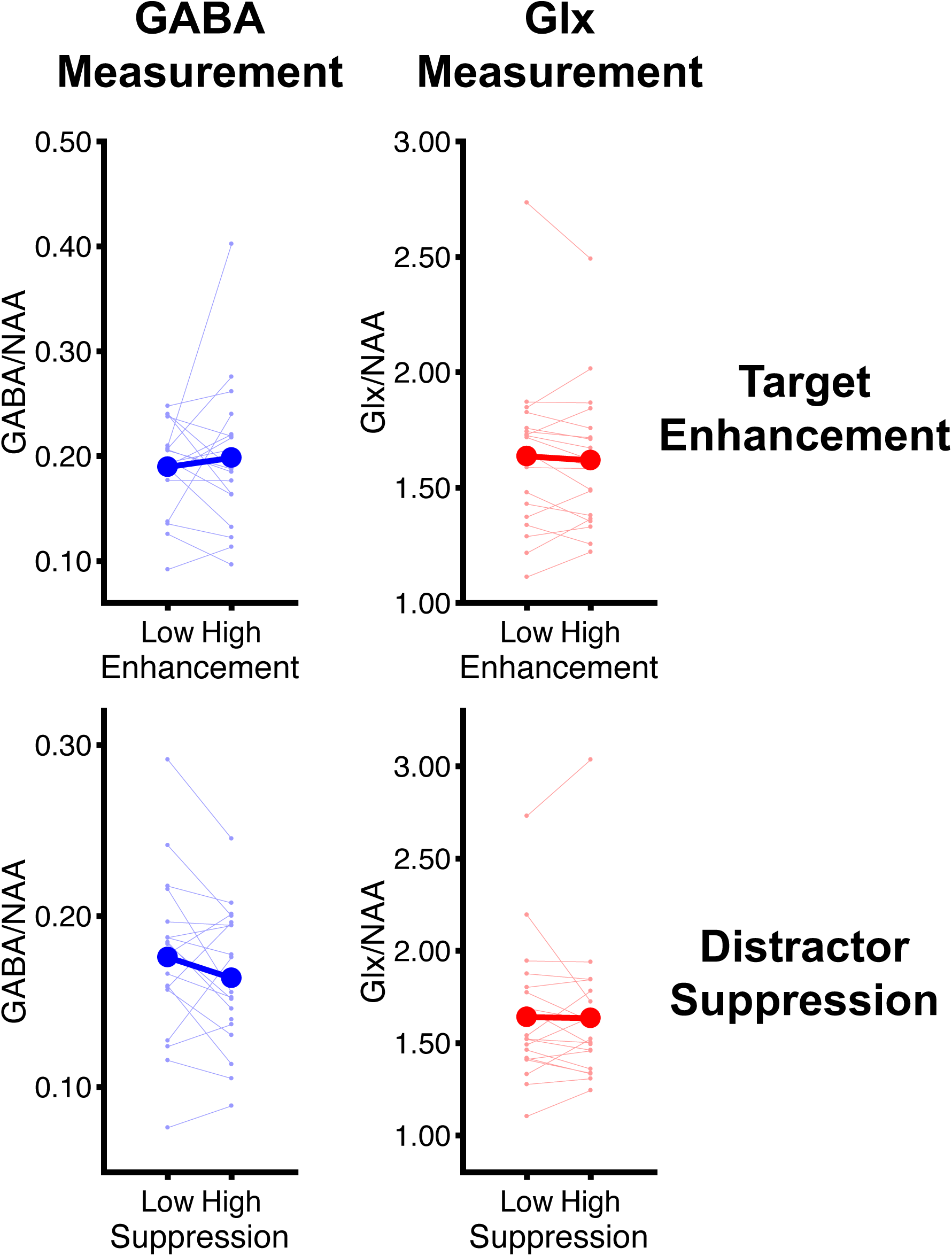
GABA and Glx measurements in the parietal VOI in the control imaging experiments. Otherwise same as Figure 3.

**Figure 9.**
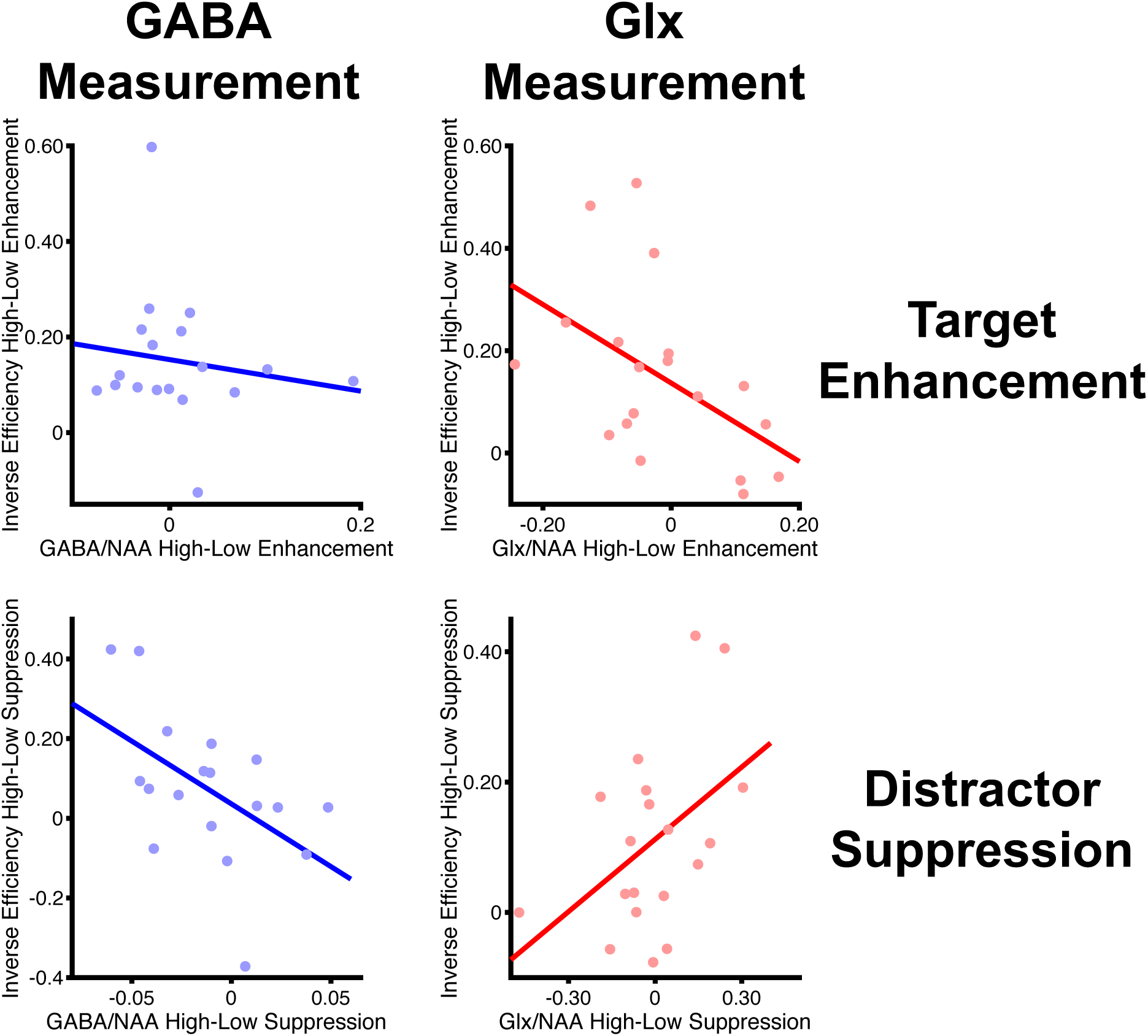
Correlational results in the control imaging experiments. More positive values on the x-axis correspond to greater concentration increase from low to high target enhancement and distractor suppression. More positive values on the y-axis correspond to greater increase in inverse efficiency score from low to high target enhancement and distractor suppression (i.e., greater decrease of response accuracy and greater increase of response time). Each correlational analysis was carried out between metabolite concentrations and tracking performance from the same fMRS run. Otherwise same as Figure 4.

## Discussion

We examined how visual signals relevant to current goals can be processed while being interfered with by goal-irrelevant distractor signals. For this purpose, we used a MOT task in which participants tracked goal-relevant targets among goal-irrelevant distractors. fMRS measurements were conducted in parietal and visual cortex while participants performed MOT with increasing tracking load to increase target-distractor interference. The results showed that GABA and Glx concentrations both increased in parietal cortex during tracking with high load compared to low load. In visual cortex only the concentration of Glx increased with load. A greater increase in GABA concentration and lower increase in Glx concentration in parietal cortex predicted better tracking performance with increasing load across participants. Control experiments showed similar correlational results between changes in parietal GABA and Glx concentrations and tracking performance when the need to suppress distractors increased during MOT. Taken together, our results suggest that GABAergic suppression of distractors in parietal cortex is critical for minimizing target-distractor interference during tracking.

Our results suggest a key role of GABAergic inhibition in parietal cortex for goal-directed visual information processing. GABA is a chief inhibitory neurotransmitter involved in refining neural responses for goal-relevant signals (De La Vega et al., 2014; Isaacson & Scanziani, 2011; Koolschijn et al., 2019; Okun & Lampl, 2008; Schmidt-Wilcke et al., 2018; H. R. Snyder et al., 2010; Tremblay et al., 2016; Tritsch et al., 2016; Wehr & Zador, 2003). The results of our main imaging experiment agree with this role of GABA as they show that GABAergic inhibition in parietal cortex increased as the need to minimize interference between targets and distractors increased in MOT with high load. Our correlational results further support the association between parietal GABA and tracking performance during pronounced target-distractor interference, as a greater increase in GABA concentration predicted better tracking performance in high load across participants. To further establish the role of GABAergic inhibition in parietal cortex for minimizing target-distractor interference, we conducted control imaging experiments in which the need to enhance targets and suppress distractors during MOT was varied separately (Bettencourt & Somers, 2009). The results showed that parietal GABA was correlated with distractor suppression, such that a greater increase in GABA concentration predicted better tracking performance when the need to suppress distractors increased. This suggests that GABA may suppress competition from spontaneous signals evoked by distractors during MOT, leading to refined processing and robust representations of goal-relevant targets in parietal cortex (Lavie, 1995; Pinsk et al., 2004; Torralbo et al., 2016). This is comparable to findings observed with other types of cognitive tasks including semantic memory and selection during language processing, which showed that increased levels of GABA sharpened the representation of relevant information for a more efficient and precise selection among competitive semantic or linguistic information (De La Vega et al., 2014; Koolschijn et al., 2019; Williams et al., 2023). It is possible that increased levels of GABA reflect a general mechanism underlying the brain’s ability to resist interference from irrelevant and unwanted signals and is universally used as a mechanism in different types of cognitive tasks (Isaacson & Scanziani, 2011; Tremblay et al., 2016; Tritsch et al., 2016). The important role of GABAergic inhibition in parietal cortex to minimize target-distractor interference is consistent with previous results from functional magnetic resonance imaging (fMRI). These previous fMRI studies showed that the representation of goal-relevant information in parietal cortex, rather than visual cortex, is resilient to interference from goal-irrelevant signals (Bettencourt & Xu, 2016; Lorenc et al., 2018; Xu, 2024).

Although our results suggest that GABAergic inhibition plays a key role in minimizing target-distractor interference, glutamatergic excitation may still be relevant to other aspects of goal-directed visual information processing. In the main imaging experiment, the concentration of Glx increased with load in parietal and visual cortex. This load-dependent increase in Glx concentration may reflect target enhancement, as our control imaging experiments showed that participants with greater increase in Glx concentration in parietal cortex also tended to perform better when the need to enhance targets during MOT increased. No such association was found between GABA concentration and target enhancement. Although we did not test this directly in our study, glutamatergic target enhancement might occur at both early and late stages of cortical visual processing as electroencephalographic recordings showed increased stimulus-evoked potentials for targets compared to distractors already in (early) visual cortex (Adamian & Andersen, 2022; Störmer et al., 2013, 2014).

Changes in GABA and Glx concentrations in parietal cortex were differentially related to changes in tracking performance with increasing load in the main imaging experiment. Higher tracking performance with increasing load was associated with greater GABA increase and lower Glx increase across participants. This suggests that excitatory processing may be most necessary when the task proves challenging to participants, whereas high-performing participants engage inhibitory processing to a greater extent. Together with the results of our control imaging experiments, this suggests that effective minimization of target-distractor interference requires GABAergic suppression of distractors to a greater extent than glutamatergic enhancement of targets.

GABA and Glx concentrations in the control imaging experiments were not significantly different between low and high target enhancement and distractor suppression conditions. Significant results were only found when changes in GABA and Glx concentrations were correlated with changes in tracking performance between conditions across participants.

Contrary to the main imaging experiment, the visual stimulation was varied between conditions in the control imaging experiments (large vs. small disks in target enhancement and a total of eight vs. twelve disks in distractor suppression), following the approach of a previous study (Bettencourt & Somers, 2009). We could speculate that significant results only in the correlational analyses indicate that changes in metabolite concentrations between conditions reflected individual differences in tracking performance rather than variations in visual stimulation.

Using fMRI, it is difficult to separate the contributions of inhibitory and excitatory processing to the blood-oxygenation-level-dependent (BOLD) response. Therefore, it is uncertain whether increased BOLD responses with increasing tracking load as reported in previous studies (Alnaes et al., 2014; Culham et al., 1998; Frank et al., 2016; Jahn et al., 2012; Jovicich et al., 2001; Maechler et al., 2025; Nummenmaa et al., 2017; Shim et al., 2010) reflect increased inhibitory or excitatory processing or a combination of both. fMRS is a novel imaging approach that overcomes this limitation of fMRI (Apšvalka et al., 2015; Craven et al., 2024; Ip & Bridge, 2022; Koolschijn et al., 2023; Lally et al., 2014; Mullins, 2018, 2024; Stanley & Raz, 2018). In addition, with fMRS it is possible to detect changes in GABA and Glx concentrations with little delay compared to fMRI (Mullins, 2018). However, fMRS also has limitations that should be considered when interpreting the results of this study. Due to the large VOI size required for fMRS, the measured concentrations reflect a combination of extracellular and intracellular neurotransmitter concentrations across large portions of the parietal and occipital lobes. Although previous studies found extended activations in the parietal and visual cortex during MOT (Adamian & Andersen, 2022; Alnaes et al., 2014; Culham et al., 2001; Frank et al., 2016; Jahn et al., 2012; Jovicich et al., 2001; Maechler et al., 2025; Nummenmaa et al., 2017; Shim et al., 2010; Störmer et al., 2013, 2014), possible differences between subregions cannot be measured with fMRS. The time-resolved fMRS design used in this study makes it likely that changes in metabolite concentrations between conditions result from neurotransmission rather than slower metabolic processes related to neurotransmitter production. However, it is still possible that both processes contributed to the measured concentration changes.

In conclusion, the results of this study suggest that GABAergic inhibition in parietal cortex is crucial for goal-directed visual information processing. By minimizing interference between target and distractors through distractor suppression, GABAergic inhibition promotes the emergence of goal-relevant target representations in parietal cortex that are robust to interference from goal-irrelevant distractors.

## Author Contribution

Z.W. and S.M.F. designed the study. Z.W., S.W., A.W., N.B., S.H., D.A., M.B., and S.M.F. collected data. Z.W. and S.M.F. analyzed data. Z.W. and S.M.F. interpreted results. Z.W. and S.M.F. wrote the first draft of the manuscript. Z.W., M.B., and S.M.F. edited and revised manuscript. S.M.F. approved final version of manuscript. S.M.F. acquired funding.

## Supplementary Material

**Supplementary Figure 1.**
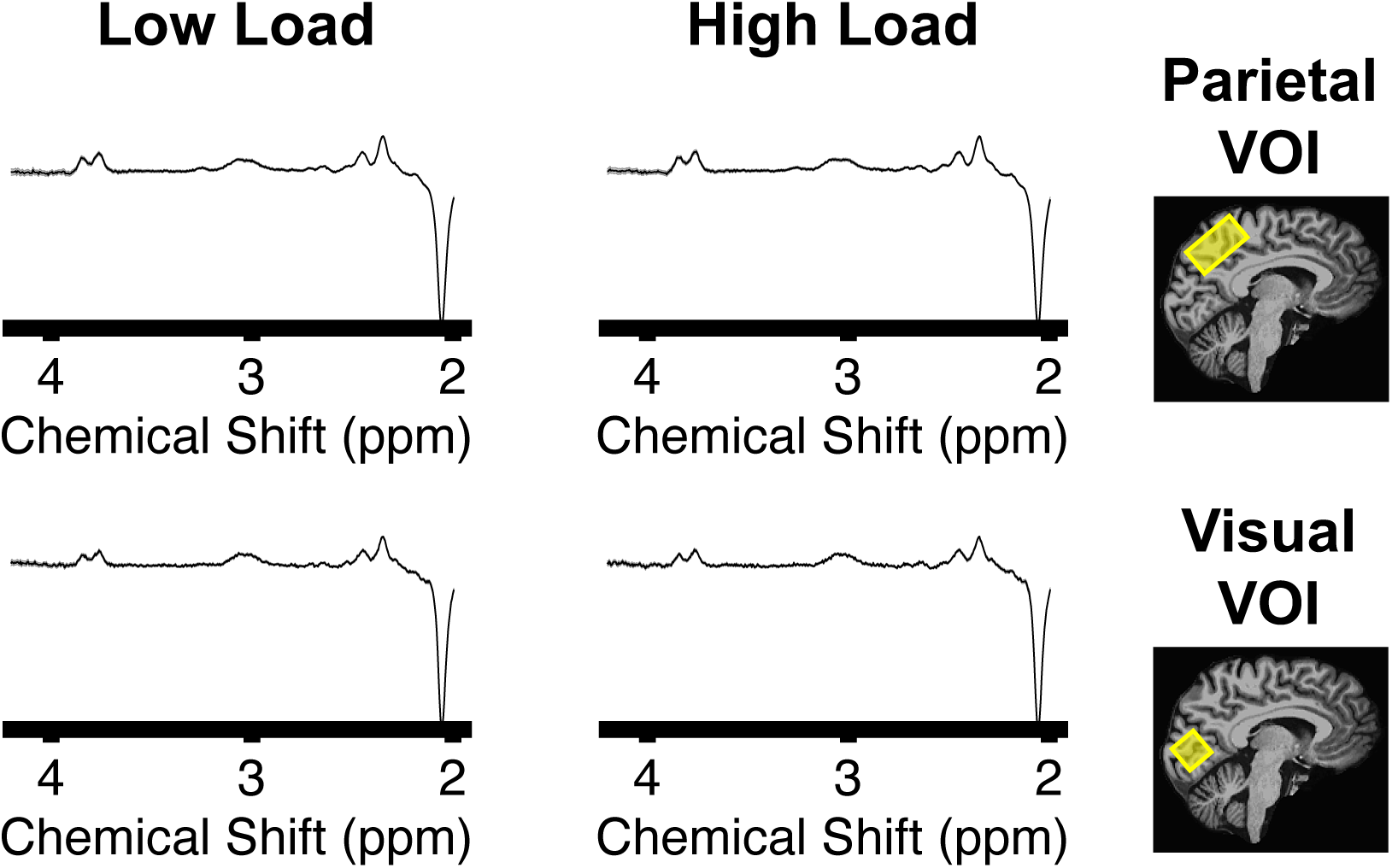
Mean MEGAPRESS difference spectra across participants in the main imaging experiment. The gray shaded area shows standard-error-of-the mean. GABA at 3 ppm and NAA at 2 ppm.

**Supplementary Figure 2.**
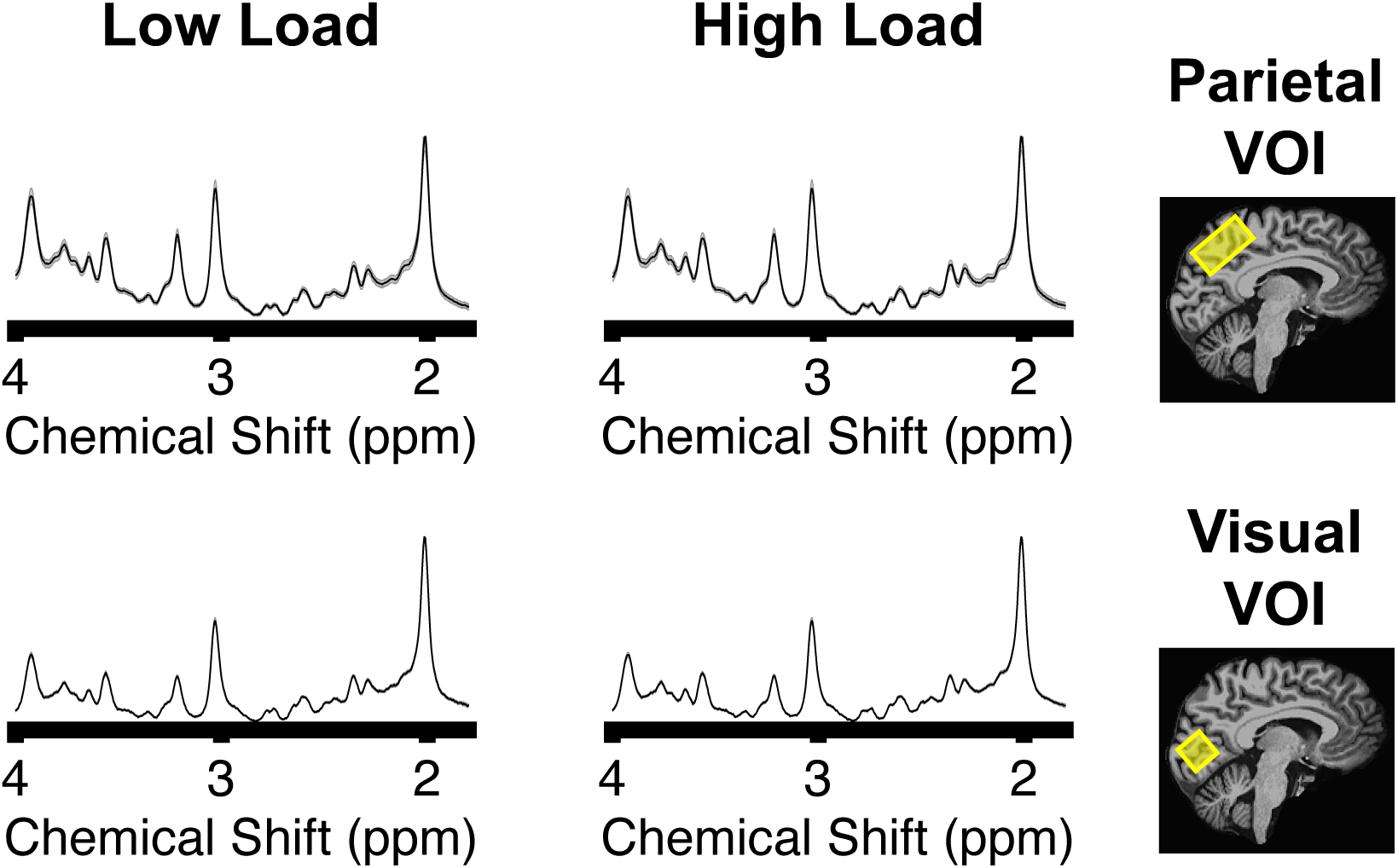
Mean PRESS spectra across participants in the main imaging experiment. Glx between 2.1 and 2.5 ppm and NAA at 2 ppm. Otherwise same as Supplementary Figure 1.

**Supplementary Figure 3.**
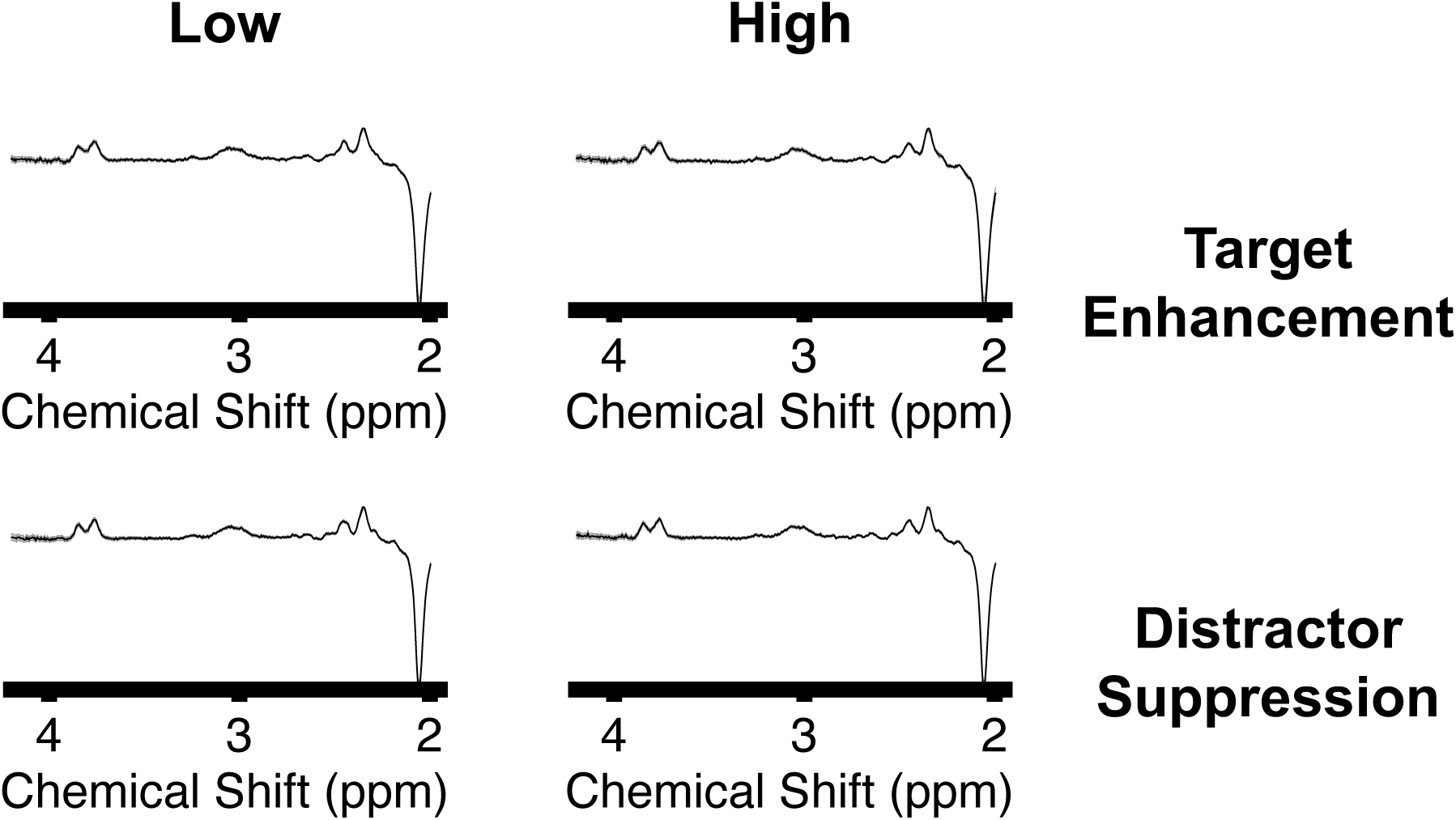
Mean MEGAPRESS difference spectra across participants in the control imaging experiments. Otherwise same as Supplementary Figure 1.

**Supplementary Figure 4.**
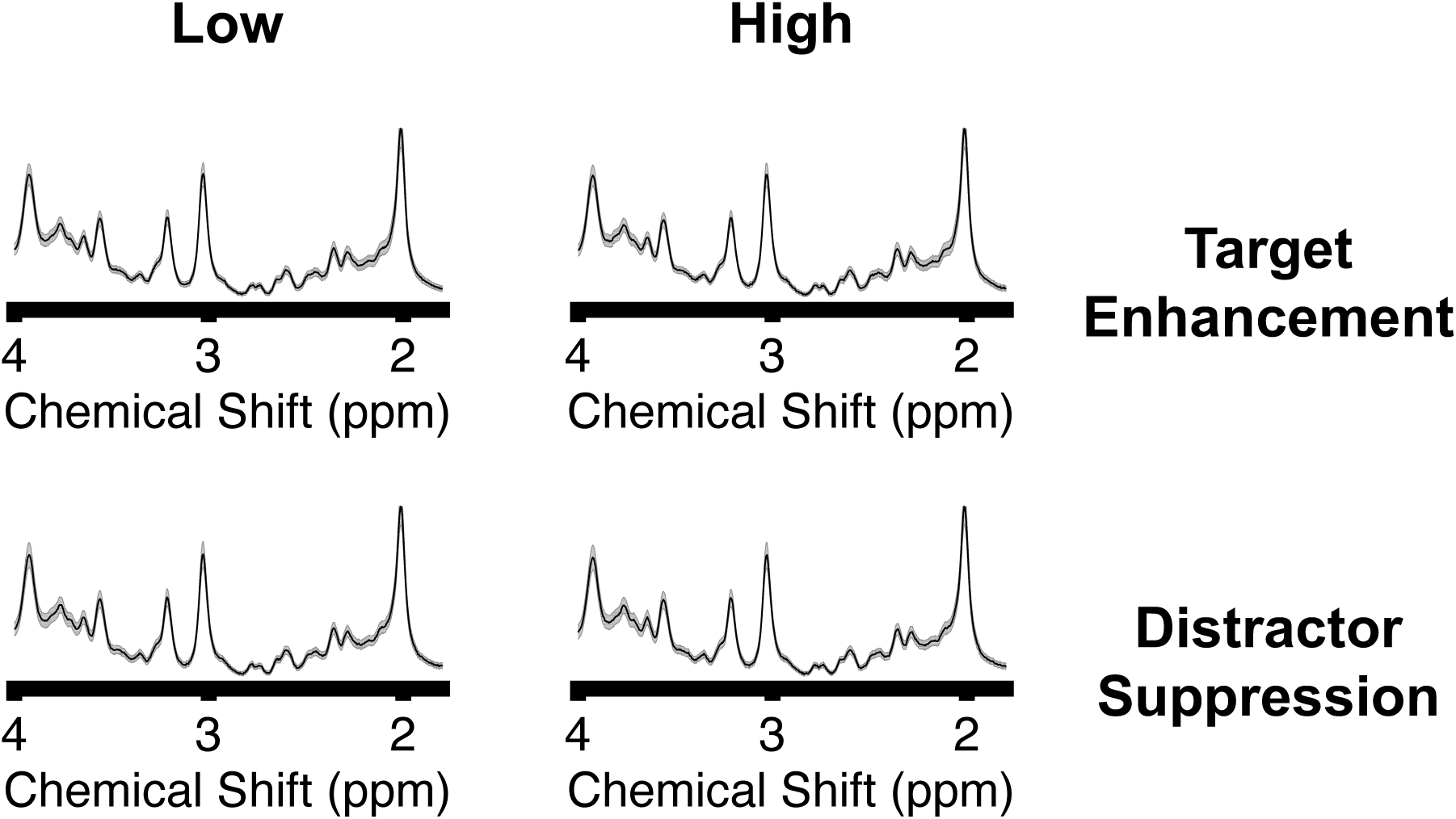
Mean PRESS spectra across participants in the control imaging experiments. Otherwise same as Supplementary Figure 2.

**Supplementary Figure 5.**
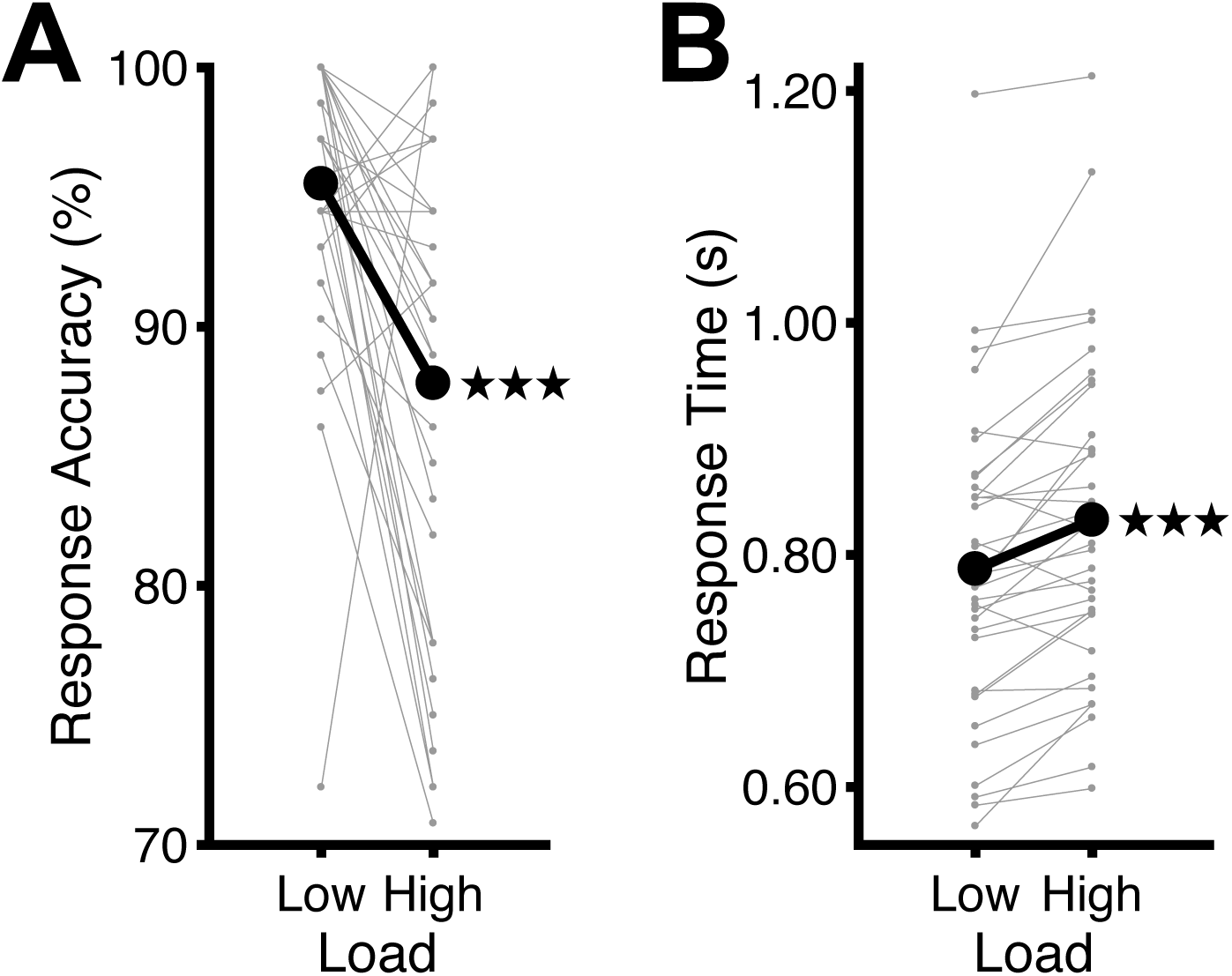
Tracking performance across fMRS runs in the main imaging experiment. (**A**) Response accuracy. Participants responded less accurately in tracking with high load than with low load [paired-sample *t*-test on arcsin-square-root transformed response accuracy; *t*(34) = −4.40, *p* < 0.001, *d* = −0.74]. Otherwise same as Figure 2. (**B**) Median response time. Participants responded slower in tracking with high load than with low load [paired-sample *t*-test on log-transformed response time; *t*(34) = 5.42, *p* < 0.001, *d* = 0.92]. Otherwise same as (A). *** *p* < 0.001.

**Supplementary Figure 6.**
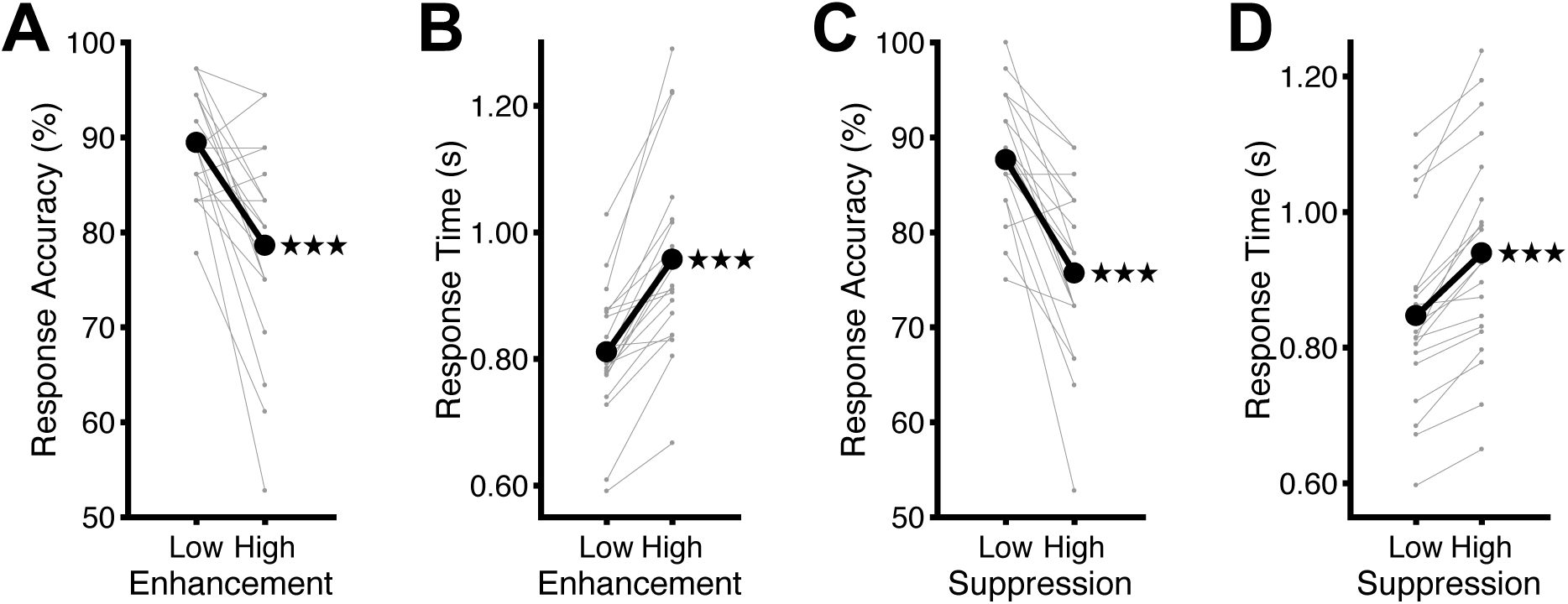
Tracking performance across runs in each control behavioral experiment. (**A**) Response accuracy in the target enhancement experiment. Participants responded less accurately in tracking with high than with low target enhancement [paired-sample *t*-test on arcsin-square-root transformed response accuracy; *t*(19) = −4.84, *p* < 0.001, *d* = −1.08]. Otherwise same as Supplementary Figure 5A. (**B**) Median response time in the target enhancement experiment. Participants responded slower in tracking with high than with low target enhancement [paired-sample *t*-test on log-transformed response time; *t*(19) = 8.24, *p* < 0.001, *d* = 1.84]. Otherwise same as Supplementary Figure 5B. (**C**) Response accuracy in the distractor suppression experiment. Participants responded less accurately in tracking with high than with low distractor suppression [paired-sample *t*-test on arcsin-square-root transformed response accuracy; *t*(19) = −6.05, *p* < 0.001, *d* = −1.35]. Otherwise same as (A). (**D**) Median response time in the distractor suppression experiment. Participants responded slower in tracking with high than with low distractor suppression [paired-sample *t*-test on log-transformed response time; *t*(19) = 8.12, *p* < 0.001, *d* = 1.82]. Otherwise same as (B). *** *p* < 0.001.

**Supplementary Figure 7.**
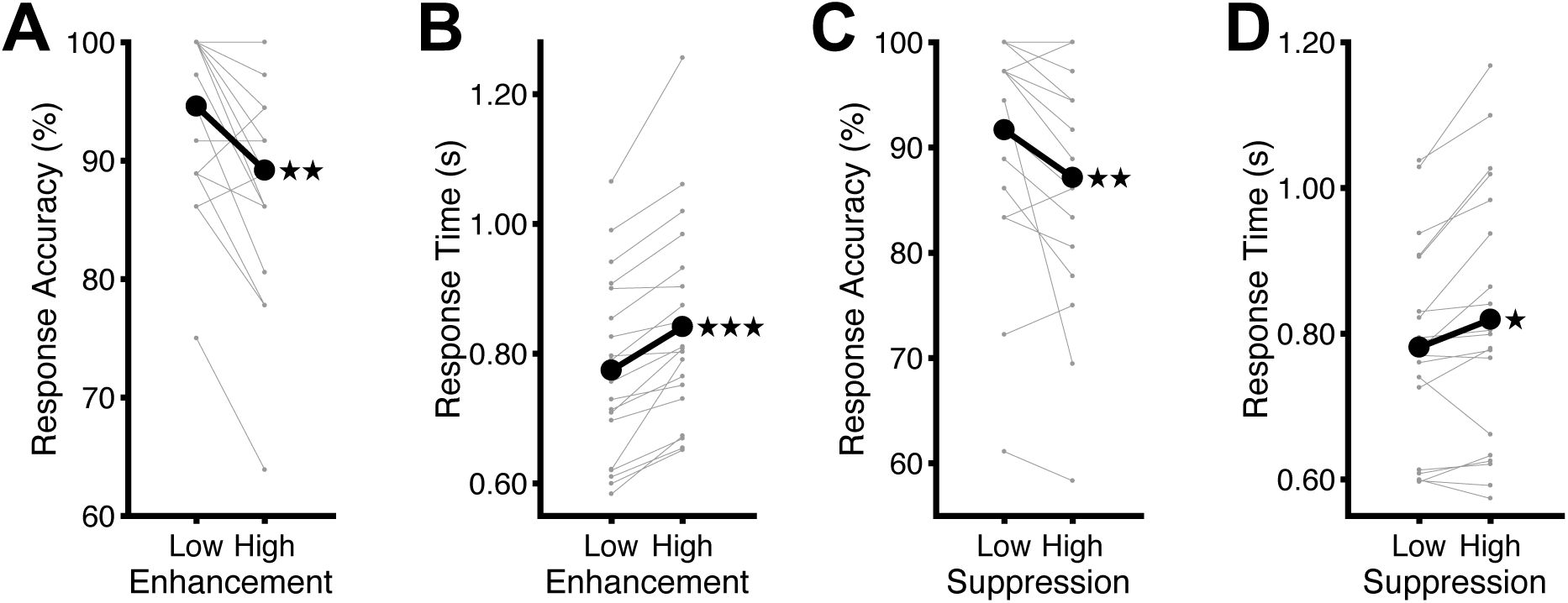
Tracking performance across fMRS runs in each control imaging experiment. (**A**) Response accuracy in the target enhancement experiment. Participants responded less accurately in tracking with high than with low target enhancement [paired-sample *t*-test on arcsin-square-root transformed response accuracy; *t*(18) = −3.89, *p* = 0.001, *d* = −0.89]. Otherwise same as Supplementary Figure 6A. (**B**) Median response time in the target enhancement experiment. Participants responded slower in tracking with high than with low target enhancement [paired-sample *t*-test on log-transformed response time; *t*(18) = 6.41, *p* < 0.001, *d* = 1.47]. Otherwise same as Supplementary Figure 6B. (**C**) Response accuracy in the distractor suppression experiment. Participants responded less accurately in tracking with high than with low distractor suppression [paired-sample *t*-test on arcsin-square-root transformed response accuracy; *t*(18) = −3.50, *p* = 0.003, *d* = −0.80]. Otherwise same as (A). (**D**) Median response time in the distractor suppression experiment. Participants responded slower in tracking with high than with low distractor suppression [paired-sample *t*-test on log-transformed response time; *t*(18) = 2.76, *p* = 0.01, *d* = 0.63]. Otherwise same as (B). * *p* < 0.05, ** *p* < 0.01, *** *p* < 0.001.

